# The effector FEP3/IRON MAN1 modulates interaction between BRUTUS-LIKE1 and bHLH subgroup IVb and IVc proteins

**DOI:** 10.1101/2021.10.07.463536

**Authors:** Daniela M. Lichtblau, Birte Schwarz, Dibin Baby, Christopher Endres, Christin Sieberg, Petra Bauer

## Abstract

Plants use the micronutrient iron (Fe) efficiently to balance the requirements for Fe during growth with its potential cytotoxic effects. A cascade of basic helix-loop-helix (bHLH) transcription factors is initiated by bHLH proteins of the subgroups IVb and IVc. This induces more than 50 genes in higher plants that can be grouped in co-expression clusters. Gene co-expression networks contain information on functional protein interactomes. We conducted a targeted yeast two-hybrid screen with pairwise combinations of 23 proteins stemming from previously characterized Fe-deficiency-induced gene co-expression clusters and regulators. We identified novel and described interactions, as well as interaction hubs with multiple interactions within the network. We found that BRUTUS-LIKE E3 ligases (BTSL1, BTSL2) interacted with basic helix-loop-helix (bHLH) transcription factors of the subgroups IVb and IVc including PYE, bHLH104 and ILR3, and with small FE UPTAKE-INDUCING PEPTIDE3/IRON MAN1 (FEP3/IMA1). Through deletion studies and with support of molecular docking, we mapped the interaction sites to three-amino-acid regions in BTSL1 and FEP3/IMA1. The FEP3/IMA1 active residues are present in interacting sites of the bHLH IVc factors. FEP3/IMA1 attenuated interaction of BTSL1 with bHLH proteins in a quantitative yeast three-hybrid assay suggesting that it is an inhibitor. Co-expression of BTSL1 and bHLH IVb and IVc factors uncovered unexpected patterns of subcellular localization. Combining deletion mapping, protein interaction and physiological analysis, we discuss the model that FEP3/IMA1 is a small effector protein inhibiting BTSL1/BTSL2-mediated degradation of bHLH subgroup IVb and IVc proteins.

**Highlights:**

- A targeted yeast two-hybrid screen of Fe deficiency-regulated proteins reveals a regulatory protein interactome consisting of E3 ligases BTS/BTSL, bHLH transcription factors of subgroups IVb and IVc and small protein FEP3/IMA1.
- Interaction sites between BTSL1, FEP3/IMA1, and bHLH IVc transcription factors were fine-mapped.
- FEP3/IMA1 is as a small effector protein that selectively attenuates the bHLH interaction with BTSL1 to regulate Fe deficiency responses.

**One sentence summary:** A targeted protein interaction screen uncovered a interactions of E3 ligase BTSL1, bHLH proteins of subgroup IVb and IVc and effector protein FEP3/IMA1 to regulate Fe deficiency responses.

## Introduction

The micronutrient iron (Fe) is a crucial cofactor for many redox and electron transfer reactions like those involved in chlorophyll synthesis and photosynthetic and respiratory electron transport chains (Nouet et al., 2011). Although very abundant in the soil, Fe is often not readily accessible for plants, because at neutral or basic pH it precipitates as insoluble Fe(III) oxides (Lindsay, 1988; Wedepohl, 1995). Fe deficiency stress results in leaf chlorosis and poor growth. To cope with low Fe availability, plants mobilize Fe in the soil, either via Fe^3+^-chelating phytosiderophores in grasses (Strategy II) or via acidification and reduction of Fe^3+^ into Fe^2+^ in non-graminaceous plants (Strategy I) (Marschner and Römheld, 1994). Once taken up, Fe is rapidly transported to local and distant sinks, or sequestered (von Wirén et al., 1999; Hell and Stephan, 2003; Briat et al., 2007; Schuler et al., 2012; Curie and Mari, 2017). Elevated cellular Fe, on the other hand, damages cellular components because of radical generation via the Fe-catalyzed Fenton reaction. Fe homeostasis is thus very critical, and Fe uptake is constantly adjusted to growth effects of hormones and environmental stress factors (Kanwar et al., 2021). For this, it is necessary that plants sense the Fe status, signal Fe demand, regulate internal Fe allocation and uptake of external Fe. The need to orchestrate these different processes is reflected in a complex transcriptomic network of co-regulated genes encoding metal ion transporters, enzymes for reduction and chelation of Fe, and transcription factors (TFs) and other regulators that steer the coordinated Fe deficiency response (Ivanov et al., 2012; Schwarz and Bauer, 2020).

Strategy I and II are controlled by a conserved cascade of basic helix-loop-helix (bHLH) TFs (Gao et al., 2019). Arabidopsis (*Arabidopsis thaliana*) has at least 12 bHLH proteins controlling Fe uptake in this cascade that are divided into subgroups according to sequences (Schwarz and Bauer, 2020). One of them, the bHLH FER-LIKE IRON DEFICIENCY INDUCED TRANSCRIPTION FACTOR (FIT) induces root Fe acquisition (Colangelo and Guerinot, 2004; Jakoby et al., 2004; Sivitz et al., 2012; Mai et al., 2016; Schwarz and Bauer, 2020). FIT targets marker genes for Fe uptake, that are *FERRIC REDUCTION OXIDASE2* (*FRO2*) encoding the Fe reductase that reduces Fe^3+^ to Fe^2+^ (Robinson et al., 1999) and *IRON-REGULATED TRANSPORTER1* (*IRT1*), which codes for the importer of Fe^2+^ (Eide et al., 1996; Vert et al., 2002). FIT acts as heterodimer together with bHLH TFs from bHLH subgroup Ib (bHLH038, bHLH039, bHLH100, bHLH101), that are together equally essential as FIT (Yuan et al., 2008; Wang et al., 2013; Trofimov et al. 2019; (Cai et al., 2021). *BHLH* subgroup Ib genes are transcriptionally induced by bHLH TFs from subgroup IVc (bHLH034, bHLH104, bHLH105 aka ILR3, bHLH115) (Zhang et al., 2015; Liang et al., 2017). Subgroup IVc bHLH TFs have redundant functions and act in a synergistic manner (Li et al., 2016; Liang et al., 2017). bHLH034, bHLH104, bHLH115 also induce transcription of BHLH subgroup IVb gene *POPEYE* (*PYE*) (Zhang et al., 2015; Liang et al., 2017), which is a direct negative regulator of Fe distribution genes *NICOTIANAMINE SYNTHASE4* (*NAS4*), *FRO3* and *ZINC-INDUCED FACILITATOR1* (*ZIF1*) (Long et al., 2010). Subgroup IVc bHLH TFs form heterodimers with the bHLH TF UPSTREAM REGULATOR OF IRT1 (URI) belonging like PYE to subgroup IVb to activate various Fe-responsive genes, including *PYE* and *BHLH* Ib TFs (Gao et al., 2019; Kim et al., 2019). The third member of the bHLH subgroup IVb family, bHLH011, in contrast, acts as a negative regulator of the Fe uptake machinery (Tanabe et al., 2019; Li et al., 2020). Despite of genetic redundancy between several bHLH subgroup IVb and IVc TFs, there are several individual functions. However, many of them are still to be uncovered.

bHLH subgroup IVc protein levels are likely controlled through proteasomal degradation. bHLH115 is targeted for degradation by BRUTUS (BTS), an Fe deficiency-induced E3 ligase and negative regulator of Fe uptake (Selote et al., 2015; Liang et al., 2017; Li et al. 2021). This somewhat paradox situation of up-regulation of negative regulator *BTS* under Fe deficiency was explained by the need to have a rapid shut-down mechanism of Fe re-mobilization (Hindt et al., 2017). Another hypothesis is that BTS ensures a constant turnover of “fresh” TF (Selote et al., 2015), a mechanism known from FIT protein activity (Sivitz et al., 2011; Meiser et al. 2011). BTS has received much attention because of its interesting domain structure that indicates Fe-sensing functions. BTS has a C-terminal REALLY INTERESTING NEW GENE (RING) domain with E3 ligase activity and N-terminal hemerythrin/HHE cation-binding motif (HHE) domains for Fe^2+^-dioxygen cofactor binding (Kobayashi et al., 2013; Selote et al., 2015; Matthiadis and Long, 2016). This structure resembles the mammalian Fe-sensing E3 ligase complex (Kobayashi et al., 2013), which has the subunit FBXL5, that is stabilized in the presence of Fe^2+^ (Salahudeen et al., 2009). Indeed, BTS protein stability and function are coupled to Fe presence or absence (Selote et al., 2015). BTS has two homologs with comparable structure, BTS-LIKE1 (BTSL1) and BTSL2, which regulate Fe uptake in a negative manner (Hindt et al., 2017). BTS, BTSL1 and BTSL2 have partly redundant functions, but only *BTS* is expressed in roots and shoots, while *BTSL1* and *BTSL2* are root-specific (Hindt et al., 2017). *BTSL2* is tightly co-regulated with *FIT*, while *BTSL1* is most similarly co-regulated with FIT targets and Fe homeostasis factors for Fe allocation (Schwarz and Bauer, 2020). In an *in vitro* assay BTSL2 ubiquitinated FIT, and FIT degradation by BTSL2 was suggested in cell-free degradation assays (Rodriguez-Celma et al. 2019). The BTS orthologue in rice OsHRZ controls orthologues of rice bHLH subgroup IVc factors (OsbHLH57/58/59/60) in a similar manner as BTS in Arabidopsis (Kobayashi, 2019). Hence, BTS and BTSL proteins act in a feedback loop to down-regulate bHLH factors for Fe homeostasis. However, this model of protein interactions is not complete and requires further proofs. Just recently, BTS was suggested to ubiquitinate and degrade a small Fe-uptake-regulatory protein (Li et al. 2021).

Fe uptake and homeostasis respond to local and systemic Fe deficiency signaling (Vert et al., 2003; Kumar et al., 2017). While the form of the signal is still unknown, hormones and small molecules modulate Fe acquisition (Brumbarova et al., 2015), and their activating effect can be overruled by phloem Fe. Mobile phloem Fe in shoots seems to be a key factor for the repression of root Fe uptake (Garcia et al., 2013 and references therein; Zhai et al., 2014; Khan et al., 2018). The shoot vasculature is important for root Fe deficiency responses (García et al., 2018 and references therein). The phloem harbors signaling proteins which are transported long-distance to elicit adaptive responses in plants. An interesting class of potential phloem-mobile proteins that are also expressed in roots are FE UPTAKE-INDUCING PEPTIDE (FEP)/IRON MAN (IMA) (Grillet et al., 2018; Hirayama et al., 2018). FEP/IMA small proteins induce Fe acquisition also in rice (Kobayashi et al., 2020), suggesting a conserved mechanism of action. FEP3/IMA1 is induced by Fe deficiency and co-regulated with bHLH subgroup IVb and IVc target genes (Schwarz and Bauer, 2020). It is currently unclear whether FEP3/IMA1 proteins themselves are a shoot-to-root signal, or whether they trigger another kind of mobile signal which ultimately leads to up-regulation of Fe acquisition genes. The mode of action of FEP/IMA-type proteins has not been known. However, a recent report suggests interaction and degradation of one of them, FEP1/IMA3, by BTS (Li et al., 2021).

In view of the number of Fe-deficiency-co-regulated genes in just the roots alone, surprisingly little is known about their interplay at the protein level. Here, we exploited Fe deficiency-regulated co-expression gene clusters to screen for novel protein interaction complexes. We identified an interactome involving BTS/BTSL E3 ligases, bHLH subgroup IVb and IVc TFs and FEP3/IMA1. We pinpointed interaction sites and we provide evidence for a mechanistic model suggesting that FEP3/IMA1 is an effector attenuating the interaction of BTSL1, likewise BTSL2, with bHLH factors of the subgroups IVb and IVc. Our results are discussed in light of a comparable mechanism just published by Li et al. (2021) on a small protein, FEP1/IMA3, that is related to FEP3/IMA1.

## Results

### A targeted yeast two-hybrid screen with co-expressed gene functions uncovered a BTS/L-bHLH IVb and IVc TFs-FEP3 interactome

Transcriptional co-regulation is often a prerequisite and indicator for protein-protein interaction, e.g. in the case of IRT1/FRO2 (Martín-Barranco et al., 2020) and bHLH039/FIT (Yuan et al. 2008). We exploited co-expression network information from Fe deficiency transcriptomics data sets (Ivanov et al.; 2012; Schwarz and Bauer, 2020) and literature to select in total 23 candidates for a targeted Y2H-based pairwise protein interaction screen, hereafter termed targeted Y2H screen (**Table 1**). Criteria for selection of candidates were (i) unknown functions of cytosolic proteins during Fe deficiency responses (at the time the study was initiated) and (ii) known regulatory functions of Fe homeostasis in the cytosol or nucleus, including proteins from outside of this network. This choice also included enzymatic functions as suspected negative controls for the screen.

**Table 1.**
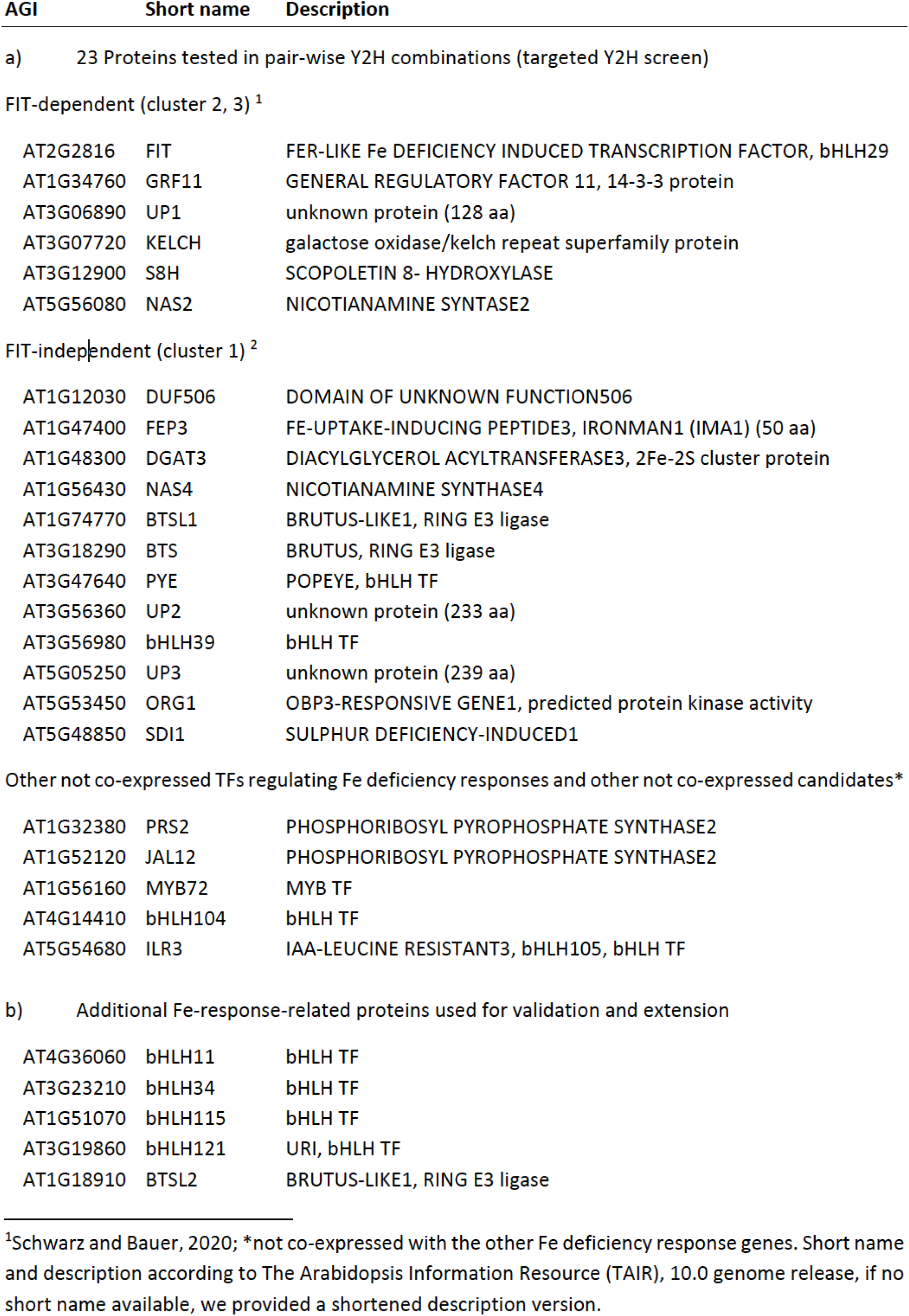
List of candidates tested in Y2H assays in this work.

At first, all 23 candidates were tested in pairwise combinations in the targeted Y2H screen (**Table 1**, **Supplemental Figure S1, S2)**. In most cases we performed reciprocal combinations and included homodimeric interaction tests. 5-6% of tested interactions were positive and they comprised 20 heterodimeric and 6 homodimeric interactions (summarized in **Figure 1**). Among them, we detected the expected interactions FIT+bHLH39, BTS+ILR3, BTS+bHLH104, PYE+ILR3 and ILR3+ILR3 (Yuan et al., 2008; Long et al., 2010; Selote et al., 2015), demonstrating the robustness of the screen. 19 heterodimer and five homodimer interactions were novel. Several interactions involved BTS or BTSL proteins and bHLH factors. For example, BTSL1 interacted with PYE, ILR3, MYB72, DUF506, SDI1, PRS2, FEP3 and UP2. Besides with BTSL1, PYE interacted with UP2 and SDI1. The targeted Y2H screen provided evidence for an Fe-regulatory interactome involving bHLH proteins of the subgroups IVb and IVc together with FEP3 and BTS/BTSL proteins. We focused on these interactions that we termed BTS/L-bHLH-FEP3 interactome. Additionally, this screen also uncovered a connection between Fe, sulfur and glucosinate metabolism via SDI interactions (Aarabi et al., 2016).

**Figure 1:**
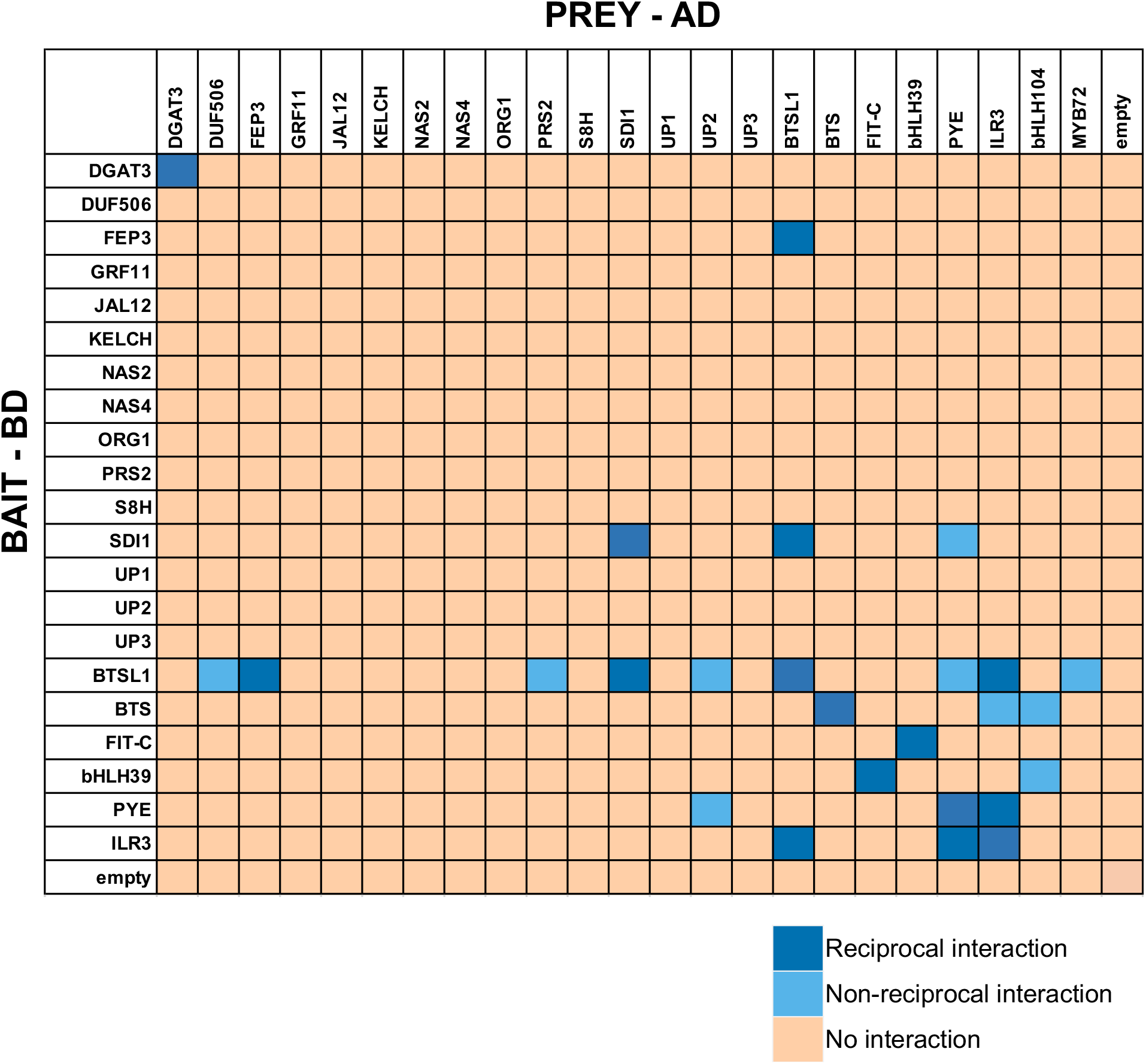
Summary results of targeted yeast two hybrid (Y2H) screen. 23 protein candidates (Table 1) were tested reciprocally in pairwise combinations in a targeted Y2H screen. Bait protein was fused to the GAL4 DNA-binding domain (BD), prey protein to the GAL4 activation domain (AD). The color code distinguishes reciprocal positive interactions (dark blue), non-reciprocal positive interactions (light blue), negative results on interactions (light orange). bHLH104 and Myb72 were not included as bait because of autoactivation. Original data are presented in Supplemental Figures S1,S2.

### The BTSL-bHLH-FEP3 interactome is validated and highlights interaction hubs that change subcellular localization of proteins

The discovered BTS/L-bHLH-FEP3 interactome attracted our attention as it suggested mechanistic insight into the functions of BTSL1, BTSL2 and FEP3 in the context of bHLH TF action. We therefore chose these proteins to enlarge and validate the targeted Y2H approach (**Figure 2, Supplemental Figure S3**). Deletion constructs were included to avoid auto-activation in the case of bHLH104-C, bHLH115-C and bHLH034-C (**Figure 2A, 2B**). BTS, BTSL1 and BTSL2 were tested against all bHLH proteins of the subgroups IVb and IVc (**Figure 2A, 2B**) and against each other (**Supplemental Figure S3A**). bHLH proteins ILR3, bHLH104-C and PYE were also tested against each other (**Supplemental Figure S3B**). This way, we found the following positive interactions: BTS interacted with URI, bHLH104-C, ILR3, bHLH115-C and bHLH034 (**Figure 2A, 2B**). BTSL1 interacted with bHLH115-C, bHLH034, ILR3, bHLH104-C, PYE and FEP3 (**Figure 2A, 2B**). BTSL2 interacted with PYE and bHLH104-C (**Figure 2A, 2B**). bHLH104-C interacted with ILR3, PYE and and homodimerized (**Supplemental Figure S3B**). Since it was reported that BTSL1 and BTSL2 interact with FIT (Rodríguez-Celma et al., 2019), we also tested specifically the interaction of BTSL1 and BTSL2 with full-length FIT and bHLH039. However, as in the targeted Y2H screen above, we did not find any proof in any combination for BTSL1 or BTSL2 interaction with FIT nor bHLH039, even though both proteins worked successfully in the other interaction pairs (**Supplemental Figure S3C, S1D**).

**Figure 2:**
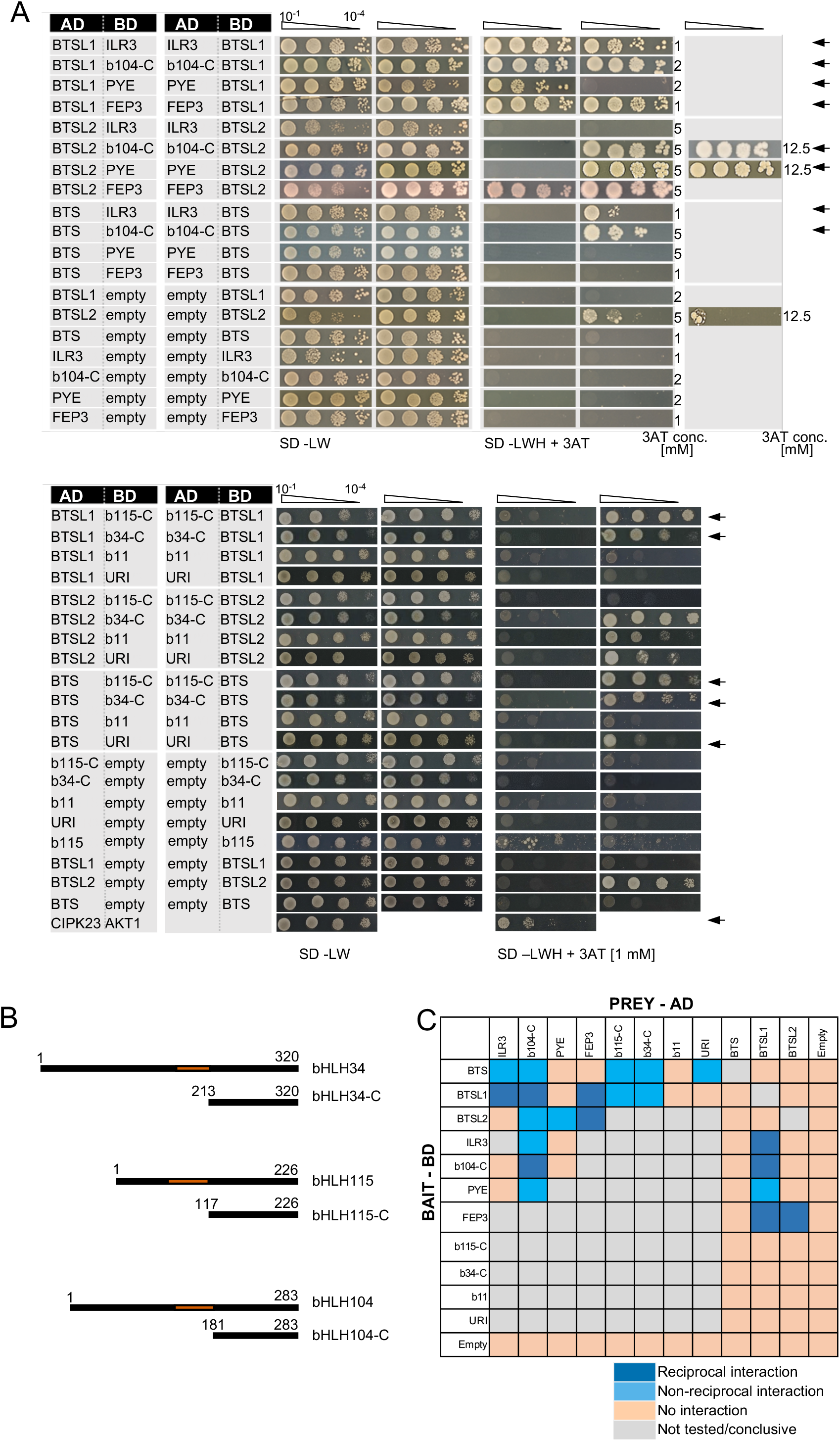
Validation of the BTS/L-bHLH-FEP3 interactome. A, BTSL1 and BTSL2 were tested in reciprocal targeted Y2H assays against various bHLH proteins of the subgroups IVb and IVc and FEP3. Yeast co-transformed with the AD and BD combinations were spotted in 10-fold dilution series (A_600_=10^-1^-10^-4^) on SD-LW (transformation control) and SD-LWH plates supplemented with different concentrations (conc.) of 3AT as indicated on the right side (selection for protein interaction). Negative controls: empty vectors. Positive control: CIPK23 and AKT1. Arrows indicate interaction. B, Schematic representation of full-length bHLH34, bHLH115, bHLH104 and their respective C-terminal parts used for Y2H. C-terminal parts lack the N-terminus and the DNA-binding domains (represented in orange). C, Summary results of A. The color code distinguishes reciprocal positive interactions (dark blue), non-reciprocal positive interactions (light blue), negative results on interactions (light orange), non-tested interactions (grey). Additional controls and Y2H validation data are presented in Supplemental Figure S3.

Taken together, the BTS/L-bHLH-FEP3 interactome was confirmed by targeted and extended Y2H data (summarized in **Figure 2C**). Some proteins within this group had a large set of interaction partners, e.g. bHLH104-C and BTSL1. Others had only one or none, e.g. URI and bHLH011. This shows specificity at the level of protein interactions. The interaction of BTSL1 and BTSL2 with FEP3 was particularly exciting since this offered the possibility of uncovering a novel mechanism of action for FEP3, BTSL1 and BTSL2. We also focused on interacting bHLH proteins ILR3, bHLH104 and PYE.

In the past, we have applied successfully bimolecular fluorescence complementation (BiFC) of YFP with simultaneous mRFP expression as control of transformation to validate protein interactions in plant cells (Gratz et all. 2019; Khan et al. 2019). When we applied this method to study BTS/BTSL protein interactions, we detected only the interaction between nY-BTSL1 and cY-PYE (**Figure 3A, top**). It was not possible to detect interactions between other fusion proteins of BTSL1 and ILR3, bHLH104 or FEP3, or any of the BTSL2 fusion proteins. In these negative cases, mRFP was detected as a control, indicating that transformation had worked (data not shown). Full-length BTSL proteins are unstable (Selote et al. 2015; Rodriguez-Selma et al. 2019). The C-terminal part of BTS-C with CHY- and CTCHY-type zinc (Zn) finger domains, a Zn ribbon domain and RING with E3 ligase function is sufficient for bHLH interaction (Selote et al., 2015). After switching to a comparable form of BTSL1-C we detected again an interaction with PYE by BiFC (**Figure 3A**, middle, nY-BTSL1-C/cY-PYE). The interaction of BTSL1 and PYE was detected in only few cells, while mRFP signals were always present in all cells of the transformed region of the leaves (**Figure 3A**). Interestingly, the YFP signals were present at the cell periphery rather than the nucleus for full-length BTSL1/PYE, while YFP signals were present in the nucleus for BTSL1-C/ PYE (**Figure 3A**, compare top and middle). Additionally, interaction of BTSL1-C and BTSL2-C with ILR3 were detected in the nucleus (**Figure 3A** bottom, nY-ILR3/cY-BTSL1-C; **Figure 3B**). No other protein interactions could be confirmed by this method despite of mRFP signals (not shown). FIT and BTSL1C were used as negative control and in none of the transformation events any positive signals were detected, while mRFP was visible (**Figure 3C**, nYFP-FIT together with cY-BTSL1-C and -BTSL2-C). Despite of the positive BiFC signals, negative data have to be carefully interpreted because of the low success rate for detecting protein interaction of BTSL1 and BTSL2 via BiFC.

**Figure 3:**
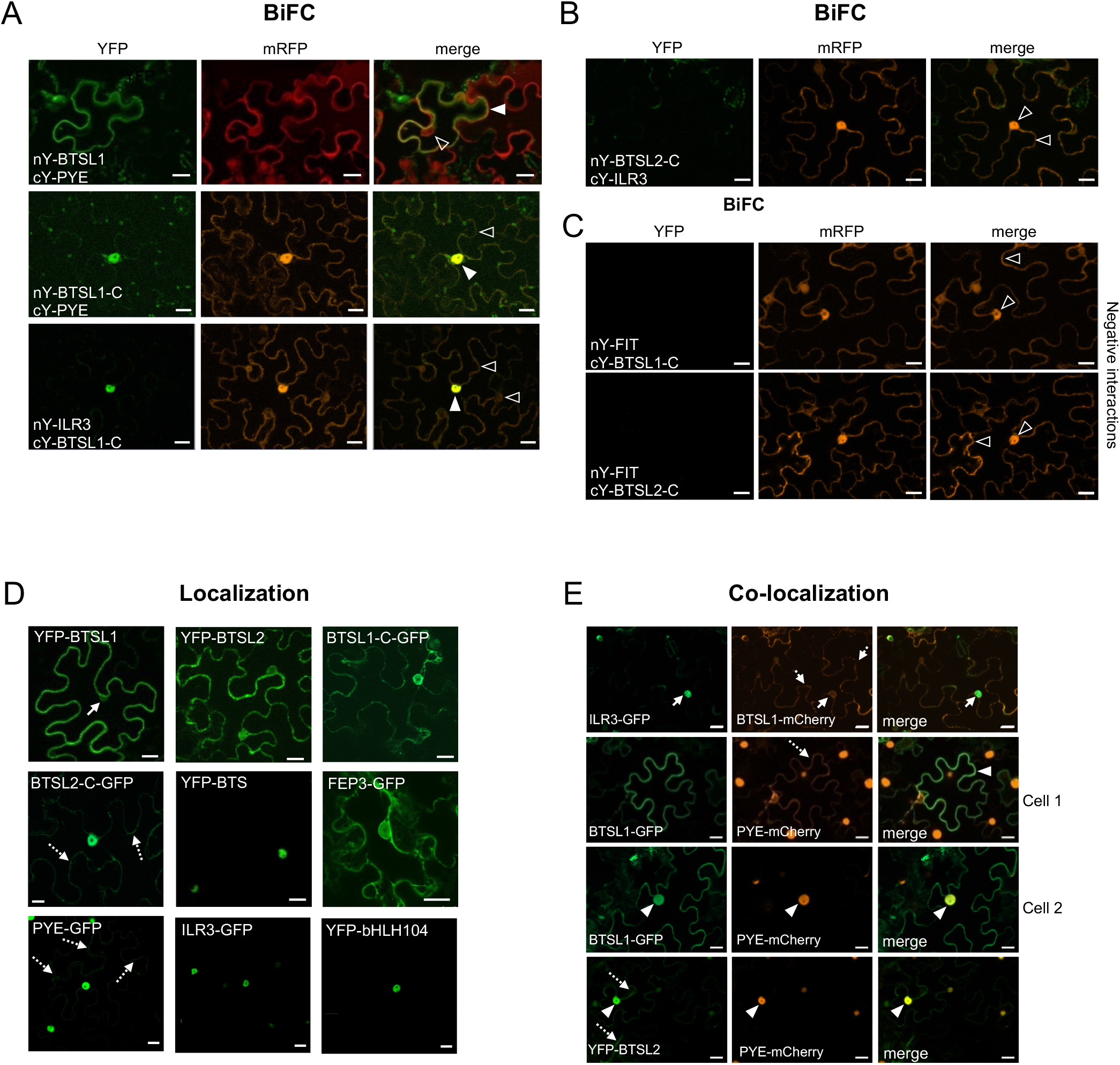
Bimolecular fluorescence complementation (BiFC) and localization/co-localization studies of the BTS/L-bHLH-FEP3 interactome. A, B, BiFC experiments showing protein interactions reflected by YFP fluorescence upon interaction of nY, nYFP- and cY, cYFP-tagged proteins, as indicated. mRFP served as positive transformation control; merge, overlay of YFP and mRFP signals. Generally, only few interactions could be demonstrated using BTSL1/BTSL1-C or BTSL2/BTSL2-C proteins, presumably because of interfering plant factors and low protein stability in plant cells. A, BTSL1 and BTSL1-C interactions with bHLH proteins. For nY-BTSL1/cY-PYE, positive YFP signals in two independent experiments with 2 infiltrated leaves of one plant, a few BiFC-positive cells per plant. For nY-BTSL1-C/cY-PYE, positive YFP signals in four independent experiments with two plants each, four to five BiFC-positive cells for each infiltrated leaf. For cY-BTSL1-C/nY-ILR3, positive signals in two independent experiments with two plants, five to ten BiFC-positive cells for each infiltrated leaf. B, BTSL2-C interaction with ILR3. C, FIT interaction with BTSL1-C and BTSL2-C. D, Subcellular localization of YFP- and GFP-tagged proteins. Respective protein fusions are indicated. For BTSL1 see confirmation of localization near the plasma membrane for BTSL1-GFP (before and after plasmolysis) and BTSL1-mCherry (Supplemental Figure S5). Non-dashed arrows indicate nucleus, dashed arrow indicates cytoplasm. E, Subcellular co-localization of GFP/YFP and mCherry-tagged proteins. Arrowheads indicate co-localization (filled arrowheads) and areas of no co-localization (un-filled arrowheads). A-E, Images were obtained after transient tobacco leaf transformation. Scale bars: 20 µm.

In summary, novel protein-protein interactions of the BTS/L-bHLH-FEP3 interactome were validated. However, in contrast to literature, no interaction was detectable for BTSL1 or BTSL2 with FIT nor bHLH039. This indicates that BTS/L interactions are specific to bHLH proteins of the subgroups IVb and IVc, fitting to data for BTS and the rice orthologs. Generally, it was difficult to study BTS/L protein interactions by BiFC in plant cells presumably because of plant factors rendering the proteins unstable, as reported previously (Selote et al., 2015).

Partners of protein interaction complexes are expected to be expressed in overlapping domains in the plant tissues and organs. To verify whether the genes encoding the BTS/L-bHLH-FEP3 interactome are co-expressed in the same root cells, we assayed promoter-driven *GUS* (β-glucuronidase) reporter gene activities in plants grown in parallel in the same growth conditions. In agreement with previous reports, the promotor activities overlapped (**Supplemental Figure S4**). *BTSL1* was expressed mainly in the outer root layers, in contrast to its proposed interaction partners *ILR3*, *BHLH104*, and *FEP3*, which are all expressed predominantly in the vascular tissue. Interestingly, the *BTSL1* expression pattern overlaps with the *PYE* expression pattern in the root differentiation zone in our analysis (**Supplemental Figure S4**). However, the location of the gene expression does not necessarily restrict the protein to the same location, as movement of PYE and FEP3 proteins was reported (Long et al. 2010; Grillet et al., 2018).

Next, we localized proteins in plant cells. As for BiFC, studies were generally hampered by the low detection of fluorescent fusions forms of proteins from the BTS/L-bHLH-FEP3 interactome. Because of that, we were only able to study intracellular localization in qualitative manner. Surprisingly, fluorescence protein-tagged BTSL1 localized mainly at the cell periphery and only weakly to the nucleus (**Figure 3D, YFP-BTSL1**). This matched the above BiFC data (see **Figure 3A**). The fluorophore position (N- /C-terminal) did not affect BTSL1 localization (**Supplemental Figure S5**). YFP-tagged BTSL2 localized to the nucleus and to the cytoplasm (**Figure 3D**, YFP-BTSL2). Remarkably, BTSL1-C-GFP and BTSL2-C-GFP localized more to the nucleus and less to the cytoplasm compared to the full-length version (**Figure 3D**, compare BTSL1-C-GFP, BTSL2-C-GFP with YFP-BTSL1 and -BTSL2), indicating an interesting pattern of BTSL1 and BTSL2 localization with an unexpected role of the N-terminal HHE domains. In contrast, YFP-BTS localized exclusively to the nucleus (**Figure 3D**, YFP-BTS). FEP3-GFP localized to nucleus and cytoplasm, with a preference of FEP3-GFP for the cytoplasm (**Figure 3D, FEP3-GFP**). PYE-GFP, ILR3-GFP and YFP-bHLH104 were located in the nucleus as expected for TFs (**Figure 3D**, PYE-GFP, ILR3-GFP, YFP-bHLH104). BTSL1-mCherry co-localized with ILR3-GFP in the nucleus, and the same was found for BTSL1-GFP and PYE-mCherry as well as YFP-BTSL2 and PYE-mCherry (**Figure 3E**). BTSL1-GFP and PYE-mCherry also co-localized outside the nucleus, whereby the PYE-mCherry signal at the cell periphery was weak compared with the nuclear signal (**Figure 3E**, compare cell1 and cell 2). It was not possible to obtain fluorescence signals for BTSL1-mCherry when it was co-expressed with FEP3-GFP (not shown). Again, protein fluorescence detection of BTSL1 and BTSL2 was hampered by low detection of signals.

In summary, these results indicate that proteins of the BTS/L-bHLH-FEP3 interactome co-localize to large extent in plant cells. Aside from the nucleus interesting dynamic cytoplasmic and cell peripheral localization effects were noted. This might, in the case of BTSL1, depend on N-terminal HHE domains.

### Interaction sites between BTSL1, bHLH proteins and FEP3 were mapped

Due to the difficulty with plant protein expression, we studied BTSL protein structure and interaction in the synthetic model yeast cell system instead. We focused on the protein interaction of BTSL1, FEP3 and the bHLH factors ILR3 and bHLH104 to dissect and fine-map protein interaction sites by Y2H, a well-established assay for these protein interaction pairs.

First, the required interaction site of BTSL1 was mapped (**Figure 4**). It was suggested that BTS interacts with its target TFs ILR3 and bHLH115 via the RING domain (Selote et al., 2015), and above BiFC data suggested the same for BTSL proteins. To pinpoint the specific interaction site, we used a deletion mutant approach to delimit further the C-terminal region of BTSL1 that is required for interaction with ILR3, bHLH104-C and FEP3 (**Figure 4**). BTSL1 and BTSL1-C interacted with bHLH factors and FEP3 (**Figure 4**, see also **Figure 2**). We divided BTSL1-C into further four deletion forms (BTSL1-C1 to -C.4, **Figure 4B**). BTSL1-C.1, lacking RING, Zn ribbon and the full CTCHY region, was not able to interact with ILR3, bHLH104-C or FEP3 (**Figure 4**). The slightly longer form BTSL1-C.2 with CHY and CTCHY domains lacked RING and Zn ribbon domains, and interacted with ILR3, bHLH104-C and FEP3 (**Figure 4**). This indicates that BTSL1 RING and Zn ribbon are not needed for this interaction. A further deletion construct BTSL1-C.3 contained only the CTCHY plus RING domains, and it interacted with ILR3 and FEP3, but not with bHLH104-C (**Figure 4**). This shows that the full CTCHY plus RING are sufficient for interaction with FEP3 and ILR3, but not for interaction with bHLH104-C. Instead, the construct BTSL1-C.4 with only RING and Zn ribbon interacted with FEP3 but none of the TF proteins (**Figure 4**). In summary, the three deletion constructs that still showed protein interaction with FEP3 (BTSL1-C.2 to -C.4) had one common small region of 14 amino acids (aa). This small 14 aa-region was named M-C site according to the first and last aa of this 14-aa stretch. M-C is located between CTCHY and RING (**Figure 4B**, yellow box). Results were less clear for ILR3 and bHLH104-C binding to BTSL1 than for FEP3 binding to BTSL1, but CTCHY and CHY domains were needed for the interaction.

**Figure 4:**
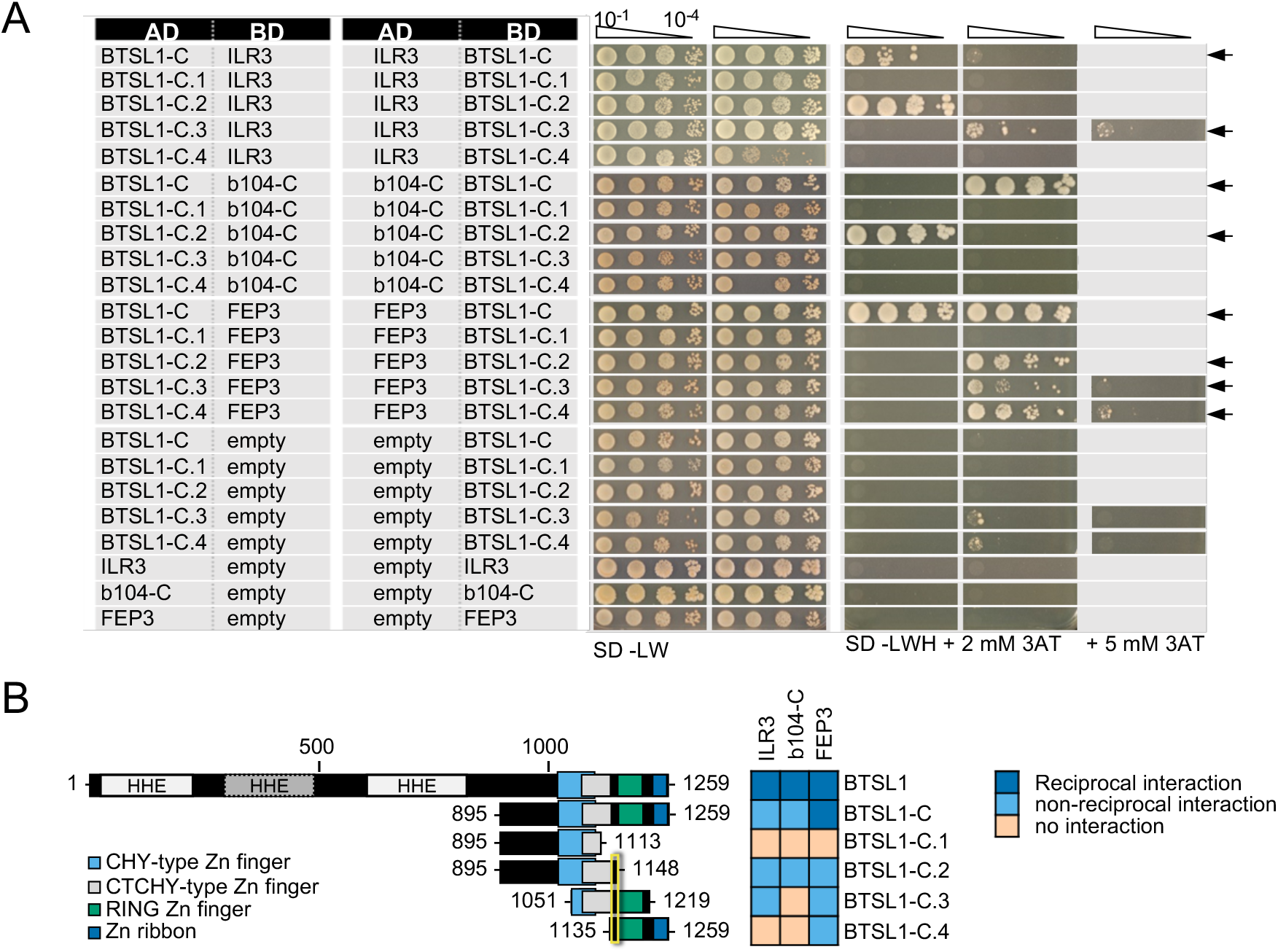
Mapping of the interaction sites in BTSL1 by yeast two hybrid (Y2H) assays. BTSL1-C and deletion forms of BTSL1-C were tested in reciprocal targeted Y2H assays against ILR3, bHLH104-C (b104-C) and FEP3. Yeast co-transformed with the AD and BD combinations were spotted in 10-fold dilution series (A_600_=10^-1^-10^-4^) on SD-LW (transformation control) and SD-LWH plates supplemented with different concentrations of 3AT as indicated (selection for protein interaction). Negative controls: empty vectors. Arrows indicate interaction. B, Schematic representation and summary of Y2H results. Left, schematic representation of full-length BTSL1, BTSL1-C and deletion constructs of BTSL1-C.1 to –C.4, used for Y2H. The domains are indicated in color. The yellow box highlights the mapped interaction site for interaction with bHLH proteins and FEP3. Right, summary results of A. The color code distinguishes reciprocal positive interactions (dark blue), non-reciprocal positive interactions (light blue), negative results on interactions (light orange).

Next, the BTSL1 M-C site was investigated in more detail and a consensus sequence within the Viridiplantae orthologs was identified (**Figure 5A**, indicated by yellow arrowhead). Deleting the M-C site in BTSL1-C (BTSL1C-dMC) abolished interactions with TFs ILR3, bHLH104-C and FEP3, indicating that the M-C site was essential (**Figure 5B, 5C**). An internal R-H part, named according to the first and last aa of an internal part, was more variable (**Figure 5C**, indicated in orange), and we found that by deleting it (BTSL1-dRH), interaction was still possible with FEP3, but not with TFs (**Figure 5B, 5C**). Substituting as a control the R-H part with a sextuple G residue spacer (BTSL1-6G) also resulted in an interaction with FEP3 but did not restore interaction with ILR3 and bHLH104-C (**Figure 5B, 5C**). Possibly, the evolutionarily conserved aa adjacent to R-H are important for interaction with FEP3 (**Figure 5B**). Thus, FEP3 interacts at a close, but separate position of BTSL1 than ILR3 and bHLH104-C. In summary, the 14-aa M-C site located close to the BTSL1 E3 RING domain is needed for interaction with FEP3, ILR3 and bHLH104-C (**Figure 5C**). Within this region, the R-H part is essential for interaction with ILR3 and bHLH104-C, but not with FEP3 (**Figure 5D**). This indicates that FEP3 and the TFs do not bind in identical manner to BTSL1.

**Figure 5:**
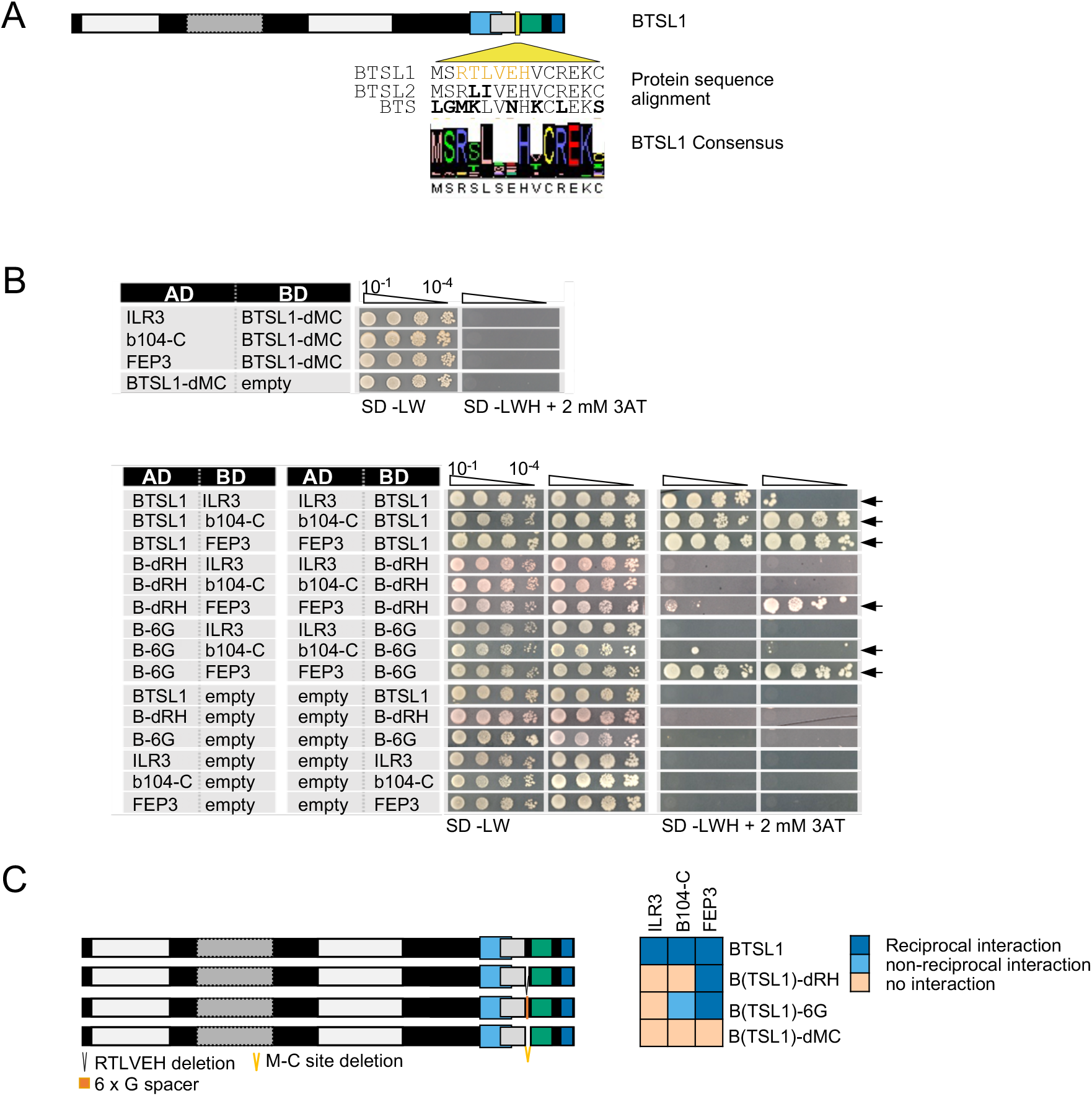
Fine-mapping of the interaction sites in BTSL1 by yeast two hybrid (Y2H) assays. A, Protein structure of BTSL1 and protein sequence alignment of the interaction M-C site within the yellow-boxed region highlighted in Figure 4B. A further sub-region is marked in orange letters, R-H site. B, Small targeted deletion forms of BTSL1-C of or within the M-C site were tested in reciprocal targeted Y2H assays against ILR3, bHLH104-C (b104-C) and FEP3. Yeast co-transformed with the AD and BD combinations were spotted in 10-fold dilution series (A_600_=10^-1^-10^-4^) on SD-LW (transformation control) and SD-LWH plates supplemented with 3AT as indicated (selection for protein interaction). Negative controls: empty vectors. Arrows indicate interaction. B, Schematic representation and summary of Y2H results. Left, schematic representation of full-length BTSL1 and deletion constructs of and within the M-C site, used for Y2H. B-6G is a sextuple glycine spacer. The regions are indicated in color. Right, summary results of B. The color code distinguishes reciprocal positive interactions (dark blue), non-reciprocal positive interactions (light blue), negative results on interactions (light orange).

Second, the interaction site within FEP3 was mapped (**Figure 6**). As shown in previous data on FEP sequence conservation across the plant kingdom (Grillet et al., 2018), FEP3 protein sequence has a conserved stretch of final 17 aa at the C-terminus (**Supplemental Figure S6**). The N-terminal half and a C-terminal half of FEP3 (termed FEP3-N and FEP3-C) were tested for their ability to interact with BTSL1, and only FEP3-C was found to be the interacting part (**Figure 6**). Next, two truncated FEP3 versions lacking the conserved stretch (FEP3-d17) or lacking the last seven aa YDYAPAA (FEP3-d7) were tested (**Figure 6**). Neither of the two constructs interacted with BTSL1, showing that the conserved stretch and the last seven aa in FEP3 are crucial for FEP3 interactions.

**Figure 6:**
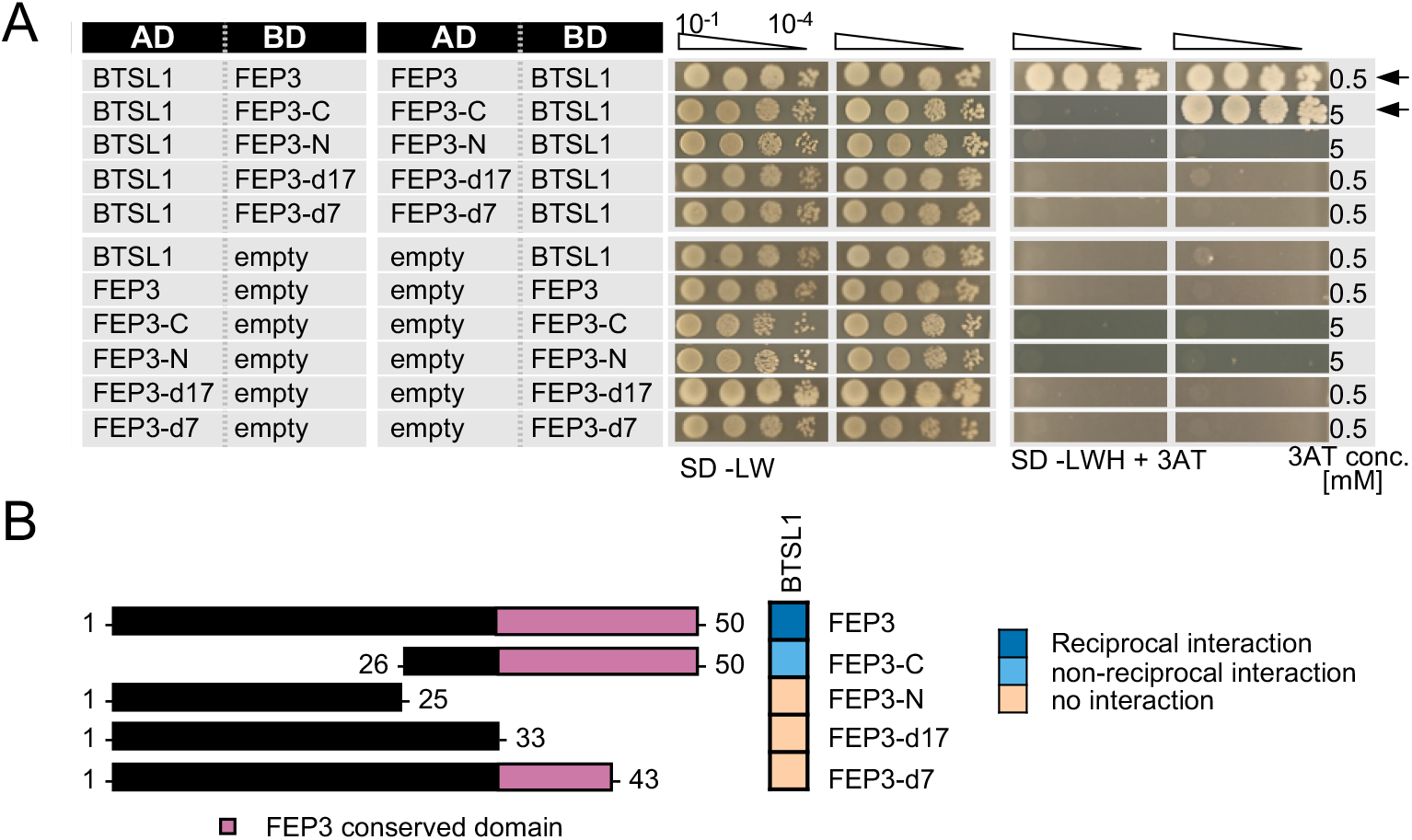
Mapping of the interaction site in FEP3 by yeast two hybrid (Y2H) assays. A, Targeted deletion forms of FEP3 were tested in reciprocal targeted Y2H assays against BTSL1. Yeast co-transformed with the AD and BD combinations were spotted in 10-fold dilution series (A_600_=10^-1^-10^-4^) on SD-LW (transformation control) and SD-LWH plates supplemented with different 3AT concentrations (conc.) as indicated (selection for protein interaction). Negative controls: empty vectors. Arrows indicate interaction. B, Schematic representation and summary of Y2H results. Left, schematic representation of FEP3 and deletion constructs, used for Y2H. Pink color illustrates the conserved region of FEP3 (see Supplemental Figure S5). Right, summary results of A. The color code distinguishes reciprocal positive interactions (dark blue), non-reciprocal positive interactions (light blue), negative results on interactions (light orange).

Third, we mapped the interaction site within the C-terminal regions of bHLH IVc proteins ILR3 and bHLH104 with BTSL1 and compared them with BTSL2 and BTS (**Figure 7**). As shown above, ILR3 interacted with BTSL1 and BTS, but not BTSL2, while bHLH104-C interacted with all three BTS/L proteins. We made an interesting observation by aligning the FEP3 sequence with the C-termini of bHLH IVc protein sequences (**Supplemental Figure S7**). We found rough similarities and conserved PAA/PVA motifs between the conserved region of FEP3 and the bHLH IVc C-termini (**Supplemental Figure S7**). In comparison, bHLH Ib protein C-termini did not align with FEP3 (data not shown). We figured that one explanation for the protein interactions could be that FEP3 mimics bHLHs IVc proteins within their last 25 aa during interaction with BTSL1/2. To test this, we constructed ILR3-d25 and bHLH104-C-d25 that lacked the 25 aa-region aligning with FEP3 YDYAPAA, and tested their ability to interact with BTS/L proteins (**Figure 7A, 7B**). We found that ILR3-d25 fragment still interacted with BTSL1, but no longer with BTS, while bHLH104-C-d25 still interacted with BTSL1 and BTSL2 but also no longer with BTS. Interestingly, short fragments only consisting of the last 25aa, ILR3-CC and bHLH104-CC, even tended to interact better with BTSL1 than the d25 fragments, while no interaction was found with BTSL2 or BTS (F**igure 7A, 7B**). Therefore, the last 25 aa of C-terminal ends of ILR3 and bHLH104 did not appear essential for interaction in all cases, but they were important. Importance of the last 25 aa was also reported in a recent study for the bHLH105 and bHLH115-BTS interaction (Li et al., 2021).

**Figure 7:**
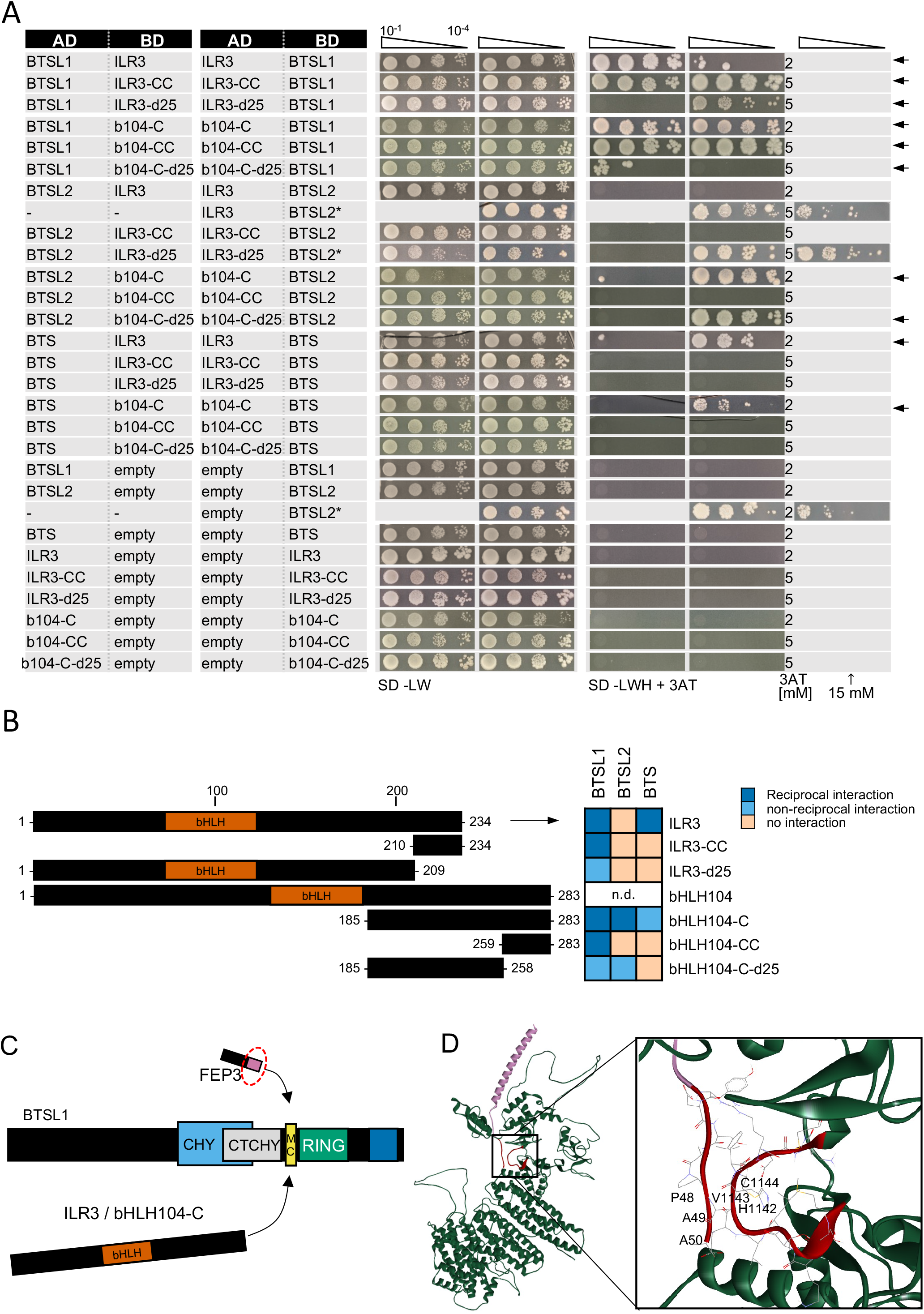
Mapping of the interaction site in bHLH subgroup IVc proteins ILR3 and bHLH104 by yeast two hybrid (Y2H) assays. A, Targeted deletion forms of bHLH proteins were tested in reciprocal targeted Y2H assays against BTSL1. Yeast co-transformed with the AD and BD combinations were spotted in 10-fold dilution series (A_600_=10^-1^-10^-4^) on SD-LW (transformation control) and SD-LWH plates supplemented with different 3AT concentrations (conc.) as indicated (selection for protein interaction). Negative controls: empty vectors. Arrows indicate interaction. B, Schematic representation and summary of Y2H results. Left, schematic representation of bHLH and deletion constructs, used for Y2H. The color illustrates the proposed region of similarity with the C-terminus of FEP3 (see Supplemental Figure S6). Right, summary results of A. The color code distinguishes reciprocal positive interactions (dark blue), non-reciprocal positive interactions (light blue), negative results on interactions (light orange). C, Proposed mechanistic model of BTSL1-C interaction at the fine-mapped M-C site with bHLH proteins of subgroup IVb and IVc and the C-terminal conserved region of FEP3. Compare with Figures 4-6 for depicted functional domains. D, Molecular homology modeling and molecular docking of BTSL1 and FEP3. Left, Homology model of BTSL1 protein predicted using AlphaFold2, used for molecular docking with FEP3. Right, details of molecular docking model between BTSL1 and FEP3. The aa highlighted are HVC within the M-C region of BTSL1 and the PAA region at the C-terminus of FEP3 that shows similarity with the C-terminus of bHLH IVb and IVc proteins. The model suggests that FEP3 is an allosteric inhibitor of bHLH binding to BTSL1.

Taken together, we were able to map interaction sites for the BTS/L-FEP3-bHLH IVc interactome (summarized in **Figure 7C**). Via AlphaFold we obtained a protein structure that we used for theoretical molecular docking experiments considering free energy values between the mapped interaction sites of BTSL1 and FEP3. Interestingly, this theoretical approach underlined experimental data and indicated precisely the three aa residues HVC within the M-C site covering with H the last aa of the R-H site of BTSL1 (**Figure 7D**). The top model that emerged indicated that PAA of last seven aa YDYAPAA of FEP3 bind to BTSL1-HVC (**Figure 7D**). As described above, PAA residues are also contained at the C-terminal end of bHLH factors (**Supplemental Figure S7**). Considering the experimental evidence, the theoretical modeling was highly convincing. Hence, through Y2H studies and molecular docking the interaction sites relevant for the BTS/L-bHLH-FEP3 interactome were fine-mapped.

### FEP3 attenuates the interaction of BTSL1 with bHLH IVb and IVc transcription factors, providing evidence for a tripartite interaction

FEP3 is a positive regulator of Fe uptake and potential phloem-mobile signal (Grillet et al., 2018). However, the mechanism by which FEP3 acts is unknown. BTSL1 and BTSL2 are suspected Fe sensors and negative regulators of Fe uptake (Hindt et al., 2017). Because FEP3 physically interacts with BTSL1 and BTSL2, and these two E3 ligases interact with bHLH IVb and IVc TFs, we hypothesized that FEP3 may inhibit BTSL1/2 to target bHLH factors. This hypothesis was strengthened by the observation, that two transgenic Arabidopsis lines over-expressing *FEP3* (FEP3-OX#1 and #3) had similar physiological phenotypes as loss-of-function defects in *btsl1 btsl2* mutants, when comparing them side-by-side (**Supplemental Figure S8,** for characterization of lines see **Supplemental Figure S9**). FEP3-Ox plants and *btsl1 btsl2* mutants had increased Fe contents per dry weight in seeds, showing that these genetic effects promote Fe accumulation (**Supplemental Figure S8A**). No difference was observed at the level of leaf chlorosis in +Fe or -Fe-exposed plants (**Supplemental Figure S8B**). However, FEP3-OX#3 and *btsl1 btsl2* mutants had longer roots at -Fe than wild type and FEP3-OX#1 (**Supplemental Figure S8C**). Increased root length is an Fe deficiency phenotype in our growth condition, dependent on active FIT (Gratz et al., 2019). These data were expected and conform with previous data on the promotion of Fe acquisition by FEP3 and an inhibitory effect of BTSL1 and BTSL2 (Grillet et al., 2018; Hindt et al., 2017). The lines showed expected and unexpected differing molecular gene expression patterns. *FEP3* and *BTSL* genes were expressed as expected in the respective overexpression and mutant situation (**Supplemental Figure S8D-8F**). However, an interesting difference was noted in that *FEP3* was not up-regulated but down-regulated in response to -Fe in *btsl1 btsl2* compared to wild type (**Supplemental Figure S8D**). Although *FEP3* is co-expressed with *BTS*, *BHLH038*, *BHLH039*, *PYE*, *FRO3* and *NAS4*, a down-regulation as seen for *FEP3* was not found in any of the co-expressed genes. Instead, with exception of non-regulated *NAS4*, all the co-expressed genes were even up-regulated at +Fe, at -Fe or in both conditions compared to wild type (**Supplemental Figure S8G-8L**). Up-regulation of *BHLH038*, *BHLH039*, *PYE*, *FRO3* was also noted in FEP3-OX lines in at least one of the Fe supply conditions, but not for *BTS* and *NAS4*, which were not regulated differently in FEP3-OX versus wild type (**Supplemental Figure S8H-L**). This suggests that BTSL proteins promote *FEP3* expression under -Fe but not expression of any of the other co-expressed genes. Unexpectedly, *ILR3* and *BHLH104* were down-regulated in FEP3-OX#1 and #3, and for *BHLH104* this was also the case in *btsl1 btsl2* under +Fe compared to the wild type (**Supplemental Figure S8M, 8N**). This was unexpected because *BHLH IVc* genes are mostly not Fe-regulated in most of our experiments as they act upstream of the Fe signaling cascade (Mai et al., 2016; Schwarz and Bauer, 2020). Thus, *ILR3* and *BHLH104* were transcriptionally regulated in an opposite manner as their downstream targets *BHLH038*, *BHLH039* and *PYE* in FEP3-OX lines. This observation was confirmed in an independent experiment using 6 d-old seedlings (**Supplemental Figure S10**). At the level of root Fe acquisition genes, *FIT* was not found differentially regulated except in FEP3-OX#3 where it was down-regulated at +Fe (**Supplemental Figure S8O**). *IRT1* and *FRO2* were co-expressed and both up-regulated in FEP3-OX#1 and in *btsl1 btsl2* at +Fe (**Supplemental Figure S8P, 8Q**), explaining the increased Fe contents.

Together, these data confirm that FEP3 acts as a positive regulator in Fe uptake, and that *btsl1 btsl2* have a similar phenotype in our growth condition as FEP3-Ox at the physiological and mostly also the molecular level. One possible explanation is that the bHLH factors ILR3 and bHLH104, which are positive regulators of bHLH subgroup Ib genes, are themselves negatively targeted by BTSL1 and BTSL2. This hypothesis predicts that FEP3 binds to BTSL1 and thereby modulates the interaction of BTSL1-bHLH subgroup IVb and IVc.

To explain the above physiology, we figured that FEP3 might inhibit BTSL1 to allow bHLH IVb and IVc TF action. This predicts that FEP3 is an effector that prevents or reduces BTSL1-bHLH interaction. We tested the effect of FEP3 on BTSL1-bHLH interactions using a quantitative yeast three hybrid (Y3H) assay. Y3H was designed to quantify ß-galactosidase activity as output of interaction between two proteins fused with AD or BD, in our case BTSL1 and a bHLH protein. Modulation of ß-galactosidase activity is quantified in the presence of an active versus inactive bridge protein, expressed under a methionine (Met)-repressible promoter, in our case FEP3 versus inactive FEP3-N. The bridge protein has either activating, repressing or neutral effect at -Met (**Figure 8A**). By testing the effect of FEP3 versus FEP3-N on FIT and bHLH39 interaction, no difference was found. Instead, high β-galactosidase values indicated strong interaction of FIT and bHLH039 in all cases irrespective of FEP3 or FEP3-N presence or absence (**Figure 8B**). This indicated that FEP3 and FEP3-N expression had a neutral effect on the FIT-bHLH039 interaction. This was expected, based on the fact that neither bHLH039 nor FIT interacted with FEP3. Interesting effects of FEP3 were seen in the case of PYE, bHLH034-C, bHLH115-C and bHLH104-C, where the presence of active FEP3 strongly impacted the protein interaction with BTSL1-C compared with the presence of inactive FEP3-N (**Figure 8C-8F**). For PYE, bHLH034-C and bHLH115-C a difference was also seen between + and -Met supporting the negative effect of FEP3. In the case of bHLH104-C no difference was seen between + and -Met, however, this interaction was also generally weaker than all other interactions, and this prevented presumably the + versus -Met effect. It must be taken into account that +Met needs to be carefully evaluated since the Met promoter tends to have leaky expression under +Met, frequently leading to a basal level expression of the bridge protein (Tirode et al., 1997). No difference occurred for BTSL1-C and ILR3 interaction in case of FEP3 or FEP3N under + and -Met (**Figure 8G**). BTSL1-C-ILR3 interaction was as strong as the one seen for FIT-C and bHLH039. Although BTSL1-C interacts with FEP3, it is possible that because of the very strong interaction, FEP3 is not able to interfere as effector with BTSL1-C and ILR3 binding.

**Figure 8:**
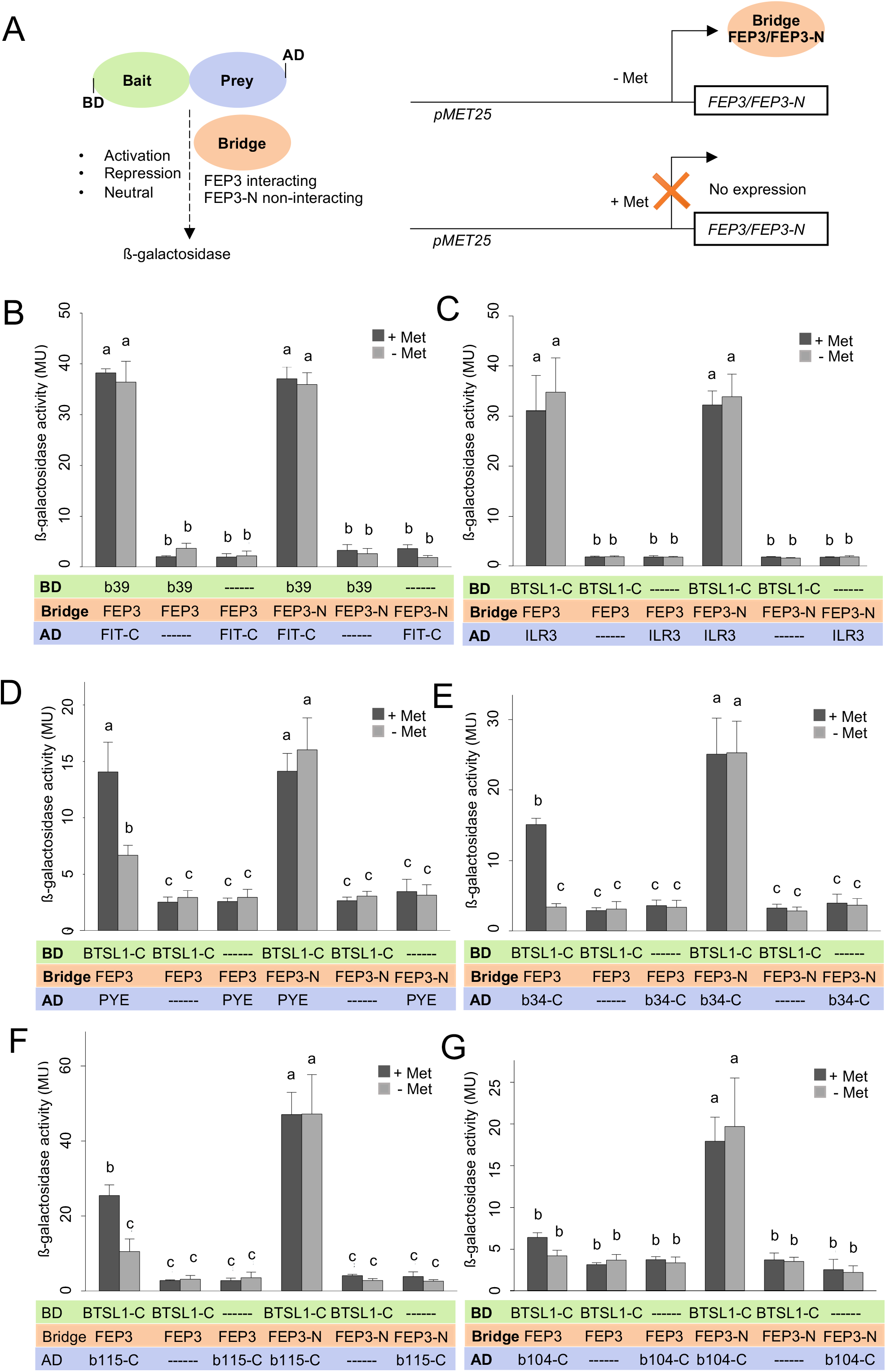
FEP3 effect on BTSL1-C and bHLH IVb and IVc interaction quantified by yeast three hybrid (Y3H) assay. A, Schematic representation of Y3H principle and design. Left, the protein interaction strength between a bait protein (fused with Gal4 DNA binding domain, BD) and prey protein (fused with Gal4 activation domain, AD) is measured by ß-galactosidase activity, here BTSL1-C and a bHLH protein (part). The effect of a bridge protein on protein interaction is measured, here interacting FEP3 and negative control non-interacting FEP-N, leading to either activation, repression or neutral effect on bait-prey protein interactions. Right, the Bridge protein is expressed under a repressible pMET25 promoter. The pMET25 promoter activity is modulated by supplementation with or without methionine (+Met, -Met). B-G, Quantification of protein interaction strengths in absence and presence of bridge protein FEP3 or FEP3-N (+/-Met) of B, FIT-C-bHLH39, C, BTSL1-C-ILR3, D, BTSL1-C-PYE, E, BTSL1-C-bHLH34-C, F, BTSL1-C-bHLH115-C, G, BTSL1-C-bHLH104-C interactions. Yeast cells are grown in SD-LWM, for bridge protein expression and SD-LW for repression of bridge protein expression; ß-galactosidase activity is determined in Miller Units (MU). Data are represented as mean values with standard deviations. Different letters indicate statistically significant differences (one-way ANOVA and Tukey’s post-hoc test, n=5, p<0.05).

Taken together, FEP3 but not FEP3-N is able to modulate the interaction of BTSL1 and bHLH proteins. FEP3 attenuates the interaction of BTSL1 with bHLH factors, provided that the interaction of BTSL1 and bHLH proteins is moderate to weak.

## Discussion

A targeted Y2H screen of 23 Fe deficiency response-related proteins revealed the protein interactome BTS/L-bHLH-FEP3/IMA1. Beyond BTS-ILR3 several other interactions between BTS/L proteins and bHLH TFs of subgroups IVb and IVc were uncovered. BTSL1 and PYE stand out as interaction hubs. FEP3/IMA1 (here named FEP3) was identified as interaction partner of BTSL1 and BTSL2. FEP3 targets via its own C-terminal end a small region termed M-C site within the C-terminus of BTSL1, conserved in BTSL2. Binding of the bHLH factors is also confined to the vicinity of this site in BTSL1. FEP3 attenuates bHLH IVb and IVc protein interaction with BTSL1, indicative of FEP3 being a small inhibitory effector protein. We propose that FEP3 competes with bHLH IVb and IVc TFs for binding sites in BTSL1 and BTSL2. This is a novel mechanism for FEP3 action. Thereby, Fe uptake and internal Fe mobilization are promoted by the action of FEP3.

### The BTS/L-bHLH subgroup IVb and IVc-FEP3 interactome is identified in the targeted Y2H screen

The targeted Y2H screen identified 14 out of 23 proteins that are candidates to interact in the context of Fe deficiency signaling, most of them were part of the combinatorial BTS/L-bHLH-FEP3 complex. The high number of protein interactions confirms the assumption that co-expressed genes encode interacting proteins, and the approach is valid to identify novel regulatory modules. Among these interactions, there were seven known interaction pairs. The fact that we found them in a targeted screen strengthens the robustness of the system. BTSL1, PYE and ILR3 occurred four or more times in different pairs of interactions including homodimers. BTS and SDI appeared at least three times including their homodimers. For this reason, we classify these proteins as “protein interaction hubs” for Fe deficiency response signaling. SDI is a sulfur response regulator (Aarabi et al., 2016), that is co-expressed with Fe deficiency genes. SDI was retrieved in this screen as a potential new interactor of BTSL1 and PYE. This information can be used to formulate new working hypotheses to test the link between sulfur, glucosinolate metabolism and Fe regulation in roots via BTSL1 and PYE and in shoots via PYE. Therefore, it can be proposed that BTSL1 and PYE have additional functions in adjusting environmental growth constraints and Fe acquisition. Our screen provides a comprehensive framework for protein interactions and evidence for a general principle that BTS/L proteins bind bHLH factors of the subgroups IVb and IVc (summarized in **Figure 9** for BTS and BTSL1). Previously, mainly bHLH115 was studied to interact with BTS (Li et al. 2021; Liang et al. 2017). The combination possibilities between bHLH-BTS/L and FEP/IMA proteins may allow cells to adequately adjust the action of the TF in a multitude of situations. In summary, the targeted Y2H screen was very valuable to detect protein interaction hubs.

**Figure 9:**
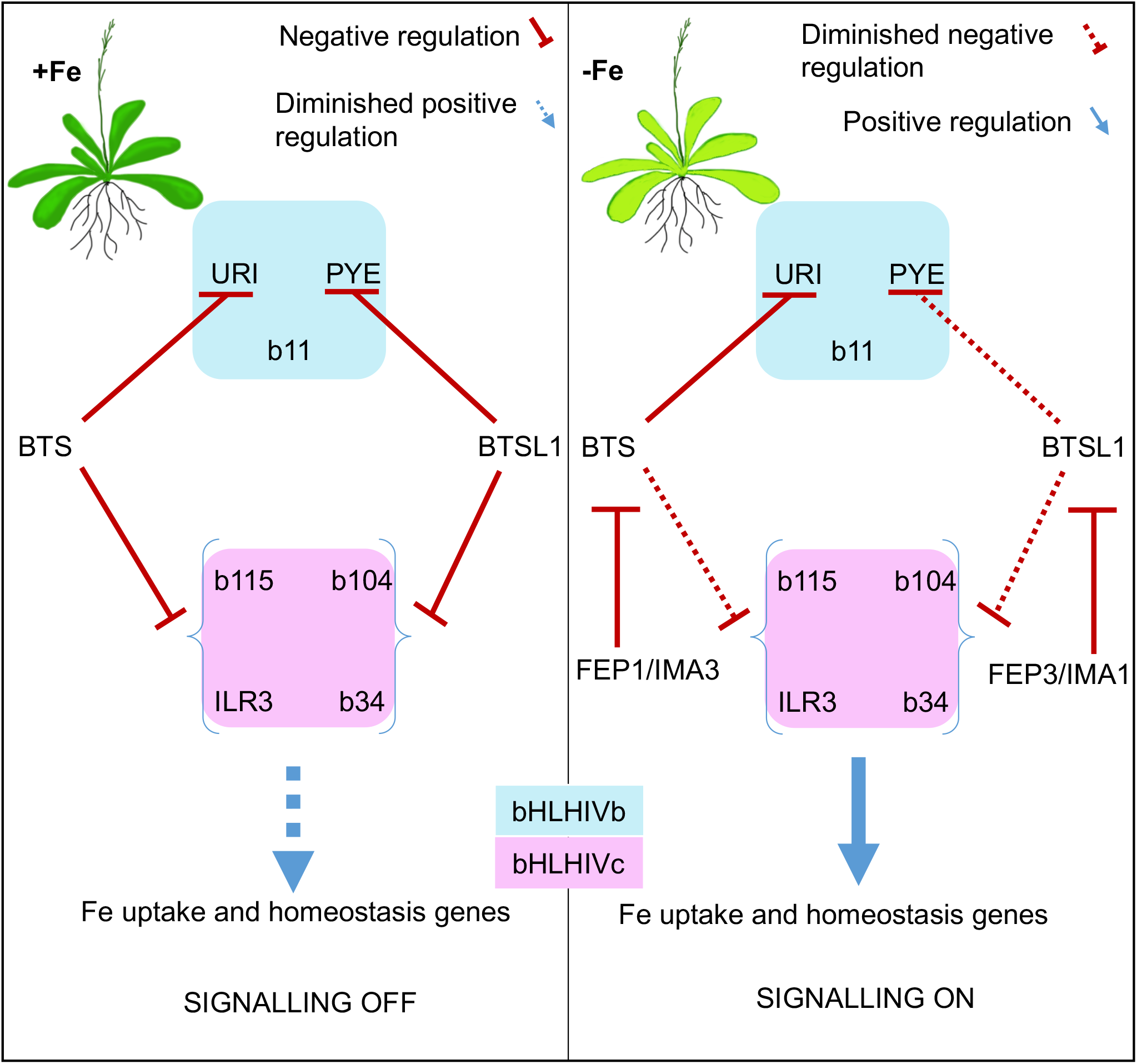
Model of FEP1 and FEP3 action to prevent BTS and BTSL1 protein-mediated degradation of bHLH factors of subgroups IVb and IVc. bHLH subgroup IVb and IVc TFs elicit Fe uptake and homeostasis in response to –Fe. These TFs are targets of E3 ligases BTS and BTSL1, possibly through degradation. BTS and BTSL1 target the same bHLH IVc but different bHLH IVb proteins. BTS, BTSL1 and the small effector proteins FEP1 and FEP3 are induced upon - Fe downstream of the bHLH IVc TFs. BTS and BTSL1 receive FEP1/FEP3 signals similar to receptor-ligand interactions. Binding of FEP1/FEP3 attenuates BTS/BTSL1-mediated degradation of bHLH TFs of subgroups IVb and IVc, allowing for enhanced Fe deficiency responses. Later, increased degradation of bHLH factors may halt Fe uptake. This model is based on intricate balancing of BTS/BTSL1-TF interaction strength and FEP1/FEP3 availability, allowing the cell to rapidly switch between on/off states to adjust Fe uptake and homeostasis. FEP3/IMA1-BTSL1-bHLH interaction was shown in this work. BTSL2 interacts with FEP3/IMA1 and with similar TF proteins of the subgroup IVb and IVc as BTSL1, and may have a similar role as BTSL1 (this work). FEP1/IMA3-BTS-bHLH interaction was shown by Li et al. (2021).

### FEP3 and bHLH subgroup IVc TFs interact through a similar C-terminal motif of BTSL1 in between CTCHY and RING domains, indicative of an inhibitor mechanism of the FEP3 effector and a tripartite interaction

E3 ligases BTS, BTSL1 and BTSL2 share a high degree of sequence similarity, suggesting that they may target identical or similar proteins. Indeed, we found that BTS and BTSL1 bind to all four members of the bHLH subgroup IVc (bHLH034, bHLH104, ILR3, bHLH115). While BTS targets subgroup IVb protein URI, BTSL1 and BTSL2 target the subgroup IVb protein PYE. This study is very complete as to the interactions of BTS/L-bHLH proteins, few data are available showing e.g. mainly BTS-ILR3 (Li et al., 2021; Liang et al. 2017). BTS/L proteins interact with FEP/IMA proteins. FEP1/IMA3 interacts with BTS (Li et al., 2021), while our study shows that FEP3/IMA1 interacts with BTSL1 and BTSL2 but not with BTS. Hence, there is specificity at the level of BTS/L interaction with bHLH factors on one side and FEP/IMA small proteins on the other side (summary in **Figure 9**). Certainly, the interactions of BTS/L, bHLH and FEP proteins diverged from a common ancestor interaction to the diversity seen in higher plants.

FEP3 is a non-secreted peptide (<100 aa) encoded by a class of evolutionarily conserved small open reading frames (ORFs) (Hsu and Benfey, 2018). Small ORF-encoded peptides (often termed SEPs) (Delcourt et al., 2018) can act in many ways, for example, as ligands to receptors or by modulating protein-protein interactions (Makarewich and Olson, 2017). As opposed to peptide hormones, SEPs do not have to be post-translationally processed in order to become bioactive peptides (Stührwohldt and Schaller, 2019). Although FEPs structurally resemble hormone peptide precursors, this and other studies could not find evidence for FEP3 or FEP1 cleavage and secretion (Grillet et al., 2018; Hirayama et al., 2018). Instead, N-terminally tagged full-length HA-FEP3 protein was detectable in plants. Therefore, FEP3 should be regarded a SEP. Interestingly, SEP-E3 ligase interactions are known from animal systems. For example, the Drosophila SEP *pri* interacts with the E3 ligase Ubr3, facilitating Ubr3 binding to the TF Svb. This changes Svb function (Zanet et al., 2015). From Drosophila as well as mammals, examples are known in which SEPs alter protein localization or bind to enzymes to affect their activity, either by direct competition with the substrate or in an allosteric manner (Cabrera-Quio et al., 2016). Here, our data suggest that FEP3 is a SEP and effector that inhibits binding of bHLH factors to an E3 ligase.

To provide evidence that FEP3 acts as SEP, we have first pinpointed the detailed interaction sites in the protein complexes with BTSL1, bHLH104/ILR3 and FEP3, and second, analyzed the effect of FEP3 on the TF-E3 ligase protein interaction complex. Very detailed deletion mapping based on Y2H interaction assays showed that FEP3 binds within the BTSL1-C terminus precisely at the M-C site which is close to the essential E3 RING domain for ubiquitination. An even smaller deletion construct with deletion of the R-H region was not suited to further delimit the FEP3 interaction. However, the interaction site was further resolved in combination with theoretical molecular docking experiments. Three aa HVC within the M-C site of BTSL1 are located right at the edge of the R-H site. HVC provide a fit with PAA at the C-terminal end of FEP3, that is essential for interaction. These same three aa PAA are also present in the required interaction region of bhLH subgroup IVc proteins. These results suggested a framework for explaining structure-function relationships and a mechanism of FEP3 action. Since BTSL1 and BTSL2 are very similar in this small region with identical functional aa residues und undergo similar protein interactions with FEP3 and bHLH subgroup IVb and IVc proteins, we predict that BTSL2 acts similarly. Likewise, the related SEP FEP1/IMA3 acts on BTS (Li et al., 2021), however, the same sequence HVC is not conserved in BTS, and also the M-C site shows multiple aa differences to the M-C site of BTSL1 and BTSL2. This explains the specificity of interaction and that FEP3 does not bind to BTS (our study). In a second step, we provided evidence for this model through quantitative assessment of the interaction strength in the presence of FEP3 in Y3H experiments. FEP3 was selective and modulated the strength of BTSL1-bHLH interactions, namely BTSL1-C-PYE, BTSL1-C-bHLH104-C, BTSL1-C-bHLH115-C and BTSL1-C-bHLH34-C. In all these cases, FEP3 caused a repression of interaction strength, suggesting competition at the BTSL1 binding site with TF fragments. In no case did we observe an increased strength of protein interaction in the presence of FEP3, excluding cooperative binding effects that stimulate the interaction. Interestingly, the very strong BTSL1-C-ILR3 interaction remained unaffected by FEP3. Thus, only the weak to moderate protein interactions between BTSL1 and bHLH TFs could be altered by FEP3, but not the strong interactions. This finding is important in the biological context to fine-tune Fe deficiency responses in balanced manner. The binding affinities of interaction partners and their intracellular concentrations must be taken into account in addition to the identities and post-translational modifications of interaction partners present in cells to calculate competition effects. These factors add an unprecedented layer of complexity to the negative regulation by the FEP3 effector mechanism.

### BTSL1 and BTS have intriguing subcellular protein localization patterns

This study showed distinct subcellular localizations of BTS, BTSL1 and BTSL2. BTSL1 was mostly located at the cell periphery and only weakly in the nucleus, in contrast to BTS, which was located only in the nucleus. BTSL2 had an intermediate localization pattern (cytoplasm and nucleus). Nuclear BTS localization was reported (Selote et al., 2015). Consistent with our results, the BTS homolog in rice, HRZ1, also localized to the nucleus while HRZ2 localized to nucleus and cytoplasm (Kobayashi et al., 2013). Because BTS, BTSL1 and BTSL2 appear to cover different subcellular compartments, they might target different parts of the Fe deficiency response cascade. For example, both BTS and its degradation target ILR3 localized and interacted in the nucleus, see also (Selote et al., 2015). When BTSL1 was expressed together with PYE or ILR3 it was localized more to the nucleus but still also at the cell periphery. This indicates that localization of BTSL1 is dependent on protein interaction partners. PYE may move from cell to cell inside the root (Long et al., 2010), and at least transiently PYE may be present in the cytoplasm or near the plasma membrane and in vicinity of plasmodesmata. It might be conceivable that upon interaction with BTSL1, the entire complex shifts to the nucleus. It might also be a possibility that another factor holds BTSL1 at the cell periphery. In this context it is interesting to note that bHLH039 is also present at the cell periphery when expressed alone. The bHLH039-FIT complex is shifted to the nucleus. bHLH039 did not interact with BTSL1, so that it is unlikely that bHLH039 is the missing link for BTSL1 localization. Interestingly, BTSL1-C was not located at the cell periphery but in cytoplasm and nucleus. We conclude that subcellular localizations of full-length BTSL1 and BTSL2 (and BTS) depend on the N-terminus and HHE domains. By binding to HHE domains, Fe^2+^ influences BTS protein stability (Selote et al., 2015), and this may likely occur also for BTSL1 and BTSL2. Because HHE domains and BTSL cytoplasmic and cell peripheral localization are linked, it is tempting to speculate that BTSL1 and BTSL2 proteins can exist in two forms, depending on the intracellular Fe status either as full-length protein mostly at the cell periphery and as truncated version that moves into the nucleus. Other E3 ligases move into the nucleus upon stress, such as AtHOS1 (HIGH EXPRESSION OF OSMOTICALLY RESPONSIVE GENES1) moving from the cytoplasm into the nucleus during cold stress (Lee et al., 2001; Dong et al., 2006) or AtRGLG1 (RING domain ligase 1) and AtRGLG2, moving from the plasma membrane into the nucleus upon ABA or salt stress treatment (Cheng et al., 2012, 2016; Belda-Palazon et al., 2019). Because HOS1, RGLG1 and RGLG2 target nuclear proteins for degradation, the nuclear-localized E3 ligases could be the active forms during the stress conditions. FEP3 was found distributed throughout the cell which is expected from a SEP. However, we were not able to localize FEP3 together with BTSL1. A reason might be that FEP3 is degraded by BTSL1, in analogy to BTS that degrades FEP1/IMA3. The cell peripheral localization of BTSL1 might also indicate a ligand-receptor-like relationship of FEP3 and BTSL1. In a hypothetical scenario, FEP3 as mobile signal binds to BTSL1 and BTSL2 in roots, which leads to de-repression of Fe acquisition. A somewhat comparable principle is known from phosphate starvation responses, where a microRNA acts as a shoot-to-root signal, targeting the mRNA of a root E2 conjugase, and thereby de-repressing the function of phosphate transporters (Huang et al., 2013).

We also tested BTS and BTSL1 promoter activities and could not confirm a clear expression partitioning in root tissue. Under -Fe *proBTS* activity was detected mostly outside the central cylinder, while *proBTS* activity under +Fe occurred mainly in the central cylinder, as reported before (Long et al. 2010). Since *proPYE* activity was detected in all root tissues, this indicates that PYE protein does not need to be cell-to-cell mobile in order to act outside the vasculature in our growth system. Growth media compositions might affect promoter gene regulation.

Taken together, the genes encoding the tripartite interactome in roots are expressed in the same or overlapping tissues of roots, additionally horizontal protein movement across the root is conceivable. BTSL1 is located at the cell periphery, while upon interaction with ILR3 and PYE the TF-E3 ligase complexes are more preponderant in the nucleus. This holds potential for the existence of a mechanism in which BTSL1 becomes truncated, switches interaction partners and relocalizes. However, it remains speculative whether BTSL1-C is a biologically relevant form of BTSL1 upon Fe deficiency stress.

### Molecular-physiological integration of the BTS/L-bHLH-FEP3 interactome

The tripartite BTSL1/BTSL2-FEP-bHLH IVb and IVc protein interactions, identified in this targeted Y2H screen, were surprising at first. Like BTS, BTSL1 and BTSL2 interacted with bHLH subgroup IVb and IVc TFs except with bHLH011. ILR3 and bHLH104 steer, along with other proteins of the bHLH subgroup IVc and URI, the top of a regulatory cascade leading to the Fe deficiency response in Arabidopsis. This points towards a role of BTS/L proteins being upstream signaling regulators, in that under Fe sufficiency conditions BTSL proteins de-regulate the Fe deficiency response pathway. A study shows that Fe deficiency is rapidly sensed in the leaves and *IMA*/*FEP* gene promoter activity is rapidly up-regulated (Khan et al., 2018). IMA/FEP proteins are reported as phloem-mobile. In a hypothetical scenario, IMA/FEP might get transported to the root system and modulate there the Fe deficiency response pathway. We could not confirm that BTSL1 or BTSL2 target FIT, and also the molecular-physiological analysis speaks against. BTSL1 and BTSL2 rather repress bHLH IVb and IVc proteins specifically in the root where *BTSL1* and *BTSL2* are expressed. This scenario is confirmed by phenotypes. When FEP3/IMA1 are overexpressed or when a *btsl1 btsl2* background is present, plants accumulate Fe in our system and in other studies of Arabidopsis (Grillet et al., 2018; Rodriguez-Celma et al., 2019; Hindt et al., 2017; Li et al., 2021). In FEP-Ox and *btsl1 btsl2*, *BHLH038*, *BHLH039*, *PYE*, *FRO3*, *IRT1* and *FRO2* were induced at +Fe in comparison with wild type. But the opposite was the case for *FEP3*, which was normally co-expressed with the other genes but now it was down-regulated in *btsl1 btsl2*. In 39Ox, high expression of only a single subgroup Ib *BHLH* gene is sufficient to cause Fe accumulation at +Fe (Naranjo Arcos et al., 2017). This leads also to a shut-down of the upstream Fe deficiency signaling cascade in 39Ox. The Fe deficiency response cascade via bHLH subgroup IVb and IVc proteins is repressed. *BTSL1* is up-regulated in 39Ox and in FEP-Ox. Therefore, up-regulation of *BTSL1* cannot be dependent on FEP3, but it is rather dependent on subgroup Ib TFs like bHLH039. Interestingly, *FIT* was up-regulated in 39Ox, but not up-regulated in FEP3-Ox, so that there may be an additional dependency on other TFs for *FIT* regulation. Clearly, the finding that *BHLH038*, *BHLH039*, *IRT1* and *FRO2* are up-regulated in *btsl1 btsl2* and in FEP Ox speaks against that FEP3 or BTSL1 and BTSL2 act at the level of FIT. Moreover, if FIT alone was a target and stabilized by the absence of BTSL1 and BTSL2 in *btsl1 btsl2* or in FEP-Ox we would have expected a phenotype of only FIT overexpression. However, FIT overexpression does not lead to Fe overaccumulation since the necessary bHLH subgroup Ib proteins are not induced along (Meiser et al., 2011). Instead, the available molecular-physiological findings rather suggest that FEP3, BTSL1 and BTSL2 act upstream of the bHLH subgroup Ib, namely to affect the regulation of bHLH subgroup IVb and IVc factors. Thereby, *BHLH038*, *BHLH039*, *IRT1* and *FRO2* are up-regulated, which explains that *btsl1 btsl2* and FEP-Ox overaccumulate Fe, similar as 39Ox. A difference is though that bHLH subgroup IVb and IVc target genes are repressed in 39Ox because of Fe sufficiency signaling. bHLH039 and other subgroup Ib proteins induce *BTSL1* and *BTSL2* expression as part of a regulatory feedback loop. In 39Ox, *BTSL1* and *BTSL2* are up-regulated and effective in repressing the Fe deficiency cascade, while this is not effective in FEP-Ox.

Interestingly, bHLH subgroup IVb and IVc TFs target genes are induced in FEP Ox and *btsl1 btsl2* indicating that Fe sufficiency-mediated signaling is not in place and not effective. PYE may positively regulate overall Fe deficiency responses, since *pye-1* mutants are chlorotic, and show e.g. decreased rhizosphere acidification and Fe reductase activity, as indicators of Fe acquisition (Long et al., 2010). PYE is a target of BTSL1 (and/or BTSL2). Perhaps, PYE also plays a role to repress *FEP3* or deregulate Fe uptake in FEP3-Ox and *btsl1 btsl2*. Li et al. (2021) proposed that TFs of the subgroups IVb or IVc repress *FEP1* gene expression, and the easiest explanation for the *FEP3* regulation is that *FEP3* is also repressed along with *FEP1* by the subgroup IVb and IVc TFs. We initially struggled to explain why *ILR3* and *BHLH104* transcripts were down-regulated in FEP3-OX and *btsl1 btsl2* mutant lines, although all their assumed downstream target genes were highly expressed. This data suggests that ILR3/bHLH104 levels are also controlled transcriptionally, possibly for a case in which the proteins are not sufficiently degraded. This might be an additional layer of control to avoid excessive Fe uptake. This scenario actually could explain why neither *btsl1 btsl2*, nor *btsl1 btsl2 bts* triple mutants (Hindt et al., 2017) or FEP3-OX plants show signs of severe Fe toxicity under +Fe. It was reported that PYE represses *ILR3* expression (Samira et al., 2018), hence *ILR3* down-regulation in FEP3-OX and *btsl1 btsl2* can be explained by the elevated *PYE* levels. In another study, ILR3 was shown to dimerize with PYE to repress *PYE* transcription (Tissot et al., 2019). We propose that IVc bHLH proteins in combination with PYE control their own transcription. Overall, though, the regulatory cascade is not fully understood yet. For example, it was expected that PYE and the genes *NAS4*, *ZIF1* and *FRO3* negatively regulated by it (Long et al., 2010) have opposite expression patterns. However, although PYE was highly expressed in mutant lines of this study, *NAS4* and *FRO3* were not down-regulated. This aligns with phenotypes of bHLH IVc gain-of-function lines (Zhang et al., 2015; Li et al., 2016), and indicates that PYE function can be bypassed or that IVc bHLHs and PYE act antagonistically to fine-tune downstream Fe acquisition and distribution genes.

### Concluding remarks

This study identified a tripartite interaction of bHLH subgroup IVb and IVc TFs competing with SEP effector protein FEP3 for BTSL1/BTSL2 E3-ligase interaction. Possibly, interacting TF proteins are targeted for degradation by BTSL1 and BTSL2 and this might be abolished by interference of FEP3. Several open questions will be of interest for future studies. BTSL1 and BTSL2 may ubiquitinate and degrade bHLH subgroup IVb and IVc TFs as a general mechanism. Since FEP3 and BTSL1 and BTSL2 have been attributed with Fe^2+^ binding abilities, Fe itself might trigger disruption of the complex and explain subcellular localization. The *FEP* family contains eight members (Grillet et al., 2018) of which *FEP1*, *FEP2* and *FEP3* are induced by -Fe in Arabidopsis (Schwarz and Bauer, 2020). FEP2 or other FEP members may also bind to BTS/BTSL proteins and exert overlapping or discrete functions with FEP1/IMA3 and FEP3/IMA1. This allows space for an intricate control through balanced combinations of individual interactions between BTS/L, bHLH and FEP/IMA proteins. Furthermore, biochemical information as to the actual affinities and concentrations of players and their post-translational modifications, e.g. as phosphorylation of URI (Kim et al., 2019), will decide on the fate of proteins.

Taken together, a mechanism in which SEPs control the function of E3 ligases as effectors through direct binding is a novel fundamental mechanism of post-translational regulatory control in plants. E3 ligases BTS/L and FEP/IMA-type small encoded peptides exert together a double negative effect. Perhaps this strong control was driven by evolutionary constraints in response to a changing environment of the plants. Identification of interaction sites with E3 ligases offers possibilities to engineer crops with modified bHLH IVb and IVc, E3 ligase or FEP/IMA binding sites. Indeed, several of the bHLH transcription factors we studied here have roles in abiotic stress protection in plants, e.g. in photoprotection (Akmakjian et al. 2021). Mechanistic understanding of bHLH subgroup IVb and IVc factors will therefore have broad impact to adapt plants to changing climate or to unravel ecological significance of Fe use efficiency during climate change.

## Materials and Methods

### Plant Material

Arabidopsis (*Arabidopsis thaliana*) ecotype Columbia-0 (Col-0) was used as wild type (WT) and as background for transgenic lines. The *btsl1 btsl2* loss-of-function double mutant was described previously (*btsl1-1 btsl2-2*, crossed SALK_015054 and SALK_048470, (Rodríguez-Celma et al., 2019). T-DNA insertion sites were verified with primer pairs LBb1.3/btsl1-1_RP (*btsl1*) and LBb1.3/btsl2-2_RP (*btsl2*) and homozygosity was verified with the primer pairs btsl1-1_LP/btsl1-1_RP and btsl2-2_LP/btsl2-2_RP (**Supplemental Table S1; Supplemental Figure 9**). For plant lines ectopically over-expressing triple HA-tagged *FEP3* (FEP3-OX) under the control of a double CaMV 35S promoter, the coding sequence was amplified from cDNA of Arabidopsis WT roots with primers carrying B1 and B2 attachment sites, respectively, transferred into the entry vector pDONR207 (Invitrogen) according to the manufacturer’s recommendations (BP reaction, Gateway, Thermo Fisher Scientific). Final constructs were obtained by transferring all candidate genes subsequently into the plant binary destination vector pAlligator2 (N-terminal triple HA fusions=HA_3_) (Bensmihen et al., 2004) via LR reactions (Thermo Fisher Scientific). Constructs were transformed into Agrobacteria (*Rhizobium radiobacter*) strain GV3101 (pMP90) (Koncz and Schell, 1986; Young et al., 2001). Stable transgenic Arabidopsis lines were generated via the Agrobacterium-mediated floral dip method (Clough and Bent, 1998). Positive transformants were selected based on seed GFP expression and genotyping PCR on the transgenic cassette, selfed and propagated to T3 generation. Insertion sites of the transgenic cassettes in FEP3-OX#1 and FEP3-OX#3 were determined by thermal asymmetric interlaced (TAIL) PCR (Liu and Whittier, 1995) with the primers S1_AL2_LB (template: gDNA), S2_AL2_LB (template: S1_AL2_LB amplicon), S3_AL2_LB (template: S2_AL2_LB amplicon), each combined with AD1, AD2, AD3, AD4, AD5, AD6. The insertion sites were verified by genotyping PCR with primer pairs FEP3-OX1_chr5 fw/S3_AL2_LB (FEP3-OX#1), FEP3-OX3_chr1 fw/S3_AL2_LB (FEP3-OX#3). Homozygosity was determined with primer pairs FEP3-OX1_chr5 fw/ FEP3-OX1_chr5 rev (FEP3-OX#1), FEP3-OX3_chr1 fw/FEP3-OX3_chr1 rev (FEP3-OX#3). Promoter sequences of *BTS* (2,994 bp), *BTSL1* (880 bp), *PYE* (1,120) and *FEP3* (1,614 bp) were amplified from Arabidopsis WT gDNA with primer pairs proBTS_-2994_B1 fw/proBTS_-2994_B2 rev (for *proBTS*), proBTSL1_-880_B1 fw/proBTSL1_-880_B2 rev (for *proBTSL1*), proPYE_-1120_B1 fw/proPYE_-1120_B2 rev (for *proPYE*), proFEP3_-1614_B1 fw/proFEP3_-1614_B2 rev (for *proFEP3*) and proFEP3_-1614_B1 fw/FEP3ns_B2 rev (for *proFEP3:FEP3*), respectively, cloned into pDONR207 (Invitrogen). Sequences were transferred into the vector pGWB3 (Nakagawa et al., 2007), generating *proBTS:GUS*, *proBTSL1:GUS*, *proPYE:GUS*, *proFEP3:GUS* and *proFEP3:FEP3-GUS* constructs. Constructs were transformed into Arabidopsis WT plants as described above. Positive transformants were selected based on hygromycin resistance and genotyping PCR, selfed and propagated to T2 or T3 generation. *ProILR3:GUS*/WT and *proBHLH104:GUS*/WT Arabidopsis lines were described.

### Plant Growth Conditions

Arabidopsis seeds were surface-sterilized and stratified. For experimental analyses seeds were distributed to sterile plates containing modified half-strength Hoagland medium (1.5 mM Ca(NO_3_)_2_, 1.25 mM KNO_3_, 0.75 mM MgSO_4_, 0.5 mM KH_2_PO_4_, 50 µM KCl, 50 µM H_3_BO_3_, 10 µM MnSO_4_, 2 µM ZnSO_4_, 1.5 µM CuSO_4_, 0.075 µM (NH_4_)_6_Mo_7_O_24_, 1% (w/v) sucrose, pH 5.8, and 1.4% w/v plant agar (Duchefa)) with (Fe sufficient, +Fe) or without (Fe deficient, -Fe) 50 μM FeNaEDTA and vertically grown in plant growth chambers (CLF Plant Climatics) under long day conditions (16h light/ 8h dark), as described in (Lingam et al., 2011). Seedlings were grown for six or ten days directly on +Fe or -Fe medium (6 day (d) system/10 d system, 6-day-old/10-day-old seedlings exposed to +/- Fe). Alternatively, seedlings were grown for 14 days on +Fe medium and then transferred for three days to either +Fe or –Fe (14+3 d system, 14-day-old plants exposed to +/- Fe), as indicated in the text.

### Yeast Two/Three-Hybrid (Y2H) Assays

#### Targeted Y2H screen

23 candidate genes were screened for interactions by pair-wise testing each as N-terminal AD (pACT2-GW constructs) and BD (pGBKT7-GW constructs) fusion proteins in both reciprocal combinations. Coding sequences were amplified from cDNA of Arabidopsis WT roots with primers carrying B1 and B2 attachment sites, respectively and transferred into pDONR207 (Invitrogen). Finally all candidate genes were transferred into destination vectors pACT2-GW and pGBKT7-GW. Yeast (*Saccharomyces cerevisiae*) strain Y187 was transformed with pACT2-GW (AD) constructs and yeast strain AH109 with pGBKT7-GW (BD) constructs via the lithium acetate (LiAc) method, based on (Gietz and Schiestl, 2007). Transformants were selected by cultivation for 2 days on minimal synthetic defined (SD) media (Clontech) lacking Leu (pACT2-GW) or Trp (pGBKT7-GW). Yeast expressing both *AD* and *BD* constructs were obtained by mating and selected on minimal SD media lacking Leu and Trp. To test for protein-protein interaction, a fresh diploid colony was resuspended in sterile H_2_O to OD_600_=1 and 10 µl of the suspensions were dropped onto minimal SD media lacking Leu, Trp and His, containing appropriate concentrations of 3-amino-1,2,4-triazole (3AT). Plates were cultivated at 30°C for up to 14 days. Diploid cells expressing each pACT2-GW:*X* construct in combination with an empty pGBKT7-GW and vice versa were used as negative controls. Combination of pGBT9.BS:*CIPK23* and pGAD.GH:*cAKT1* was used as a positive control of the system (Xu et al., 2006).

#### Targeted Y2H assays for validation

Selected protein pairs of the Y2H screen plus additional proteins (URI, bHLH11, bHLH34, bHLH115) and mutagenized/truncated protein versions were assayed as N-terminal AD and BD fusion proteins in both reciprocal combinations. Mutagenized *BTSL1* versions *BTSL1-dRH*, *BTSL1-6G and BTSL1-dMC* were created as described in “BTSL1 mutagenesis”. Truncated versions *BTSL1-N*, *BTSL1-C*, *BTSL1-C.1*, *BTSL1-C.2*, *BTSL1-C.3*, *BTSL1-C.4*, *FEP3-N*, *FEP3-C*, *FEP3-d7*, *ILR3-d25*, *ILR3-CC*, *bHLH104-C*, *bHLH104-C-d25*, *bHLH104-CC* were amplified with primers listed in **Supplemental Table S1** and cloned into pACT2-GW and pGBKT7-GW as described in the previous section. Yeast strain AH109 was co-transformed with both pACT2-GW:*X* (AD-X) and pGBKT7-GW:*Y* (BD-Y) (including empty vector controls) as described in the previous section. X and Y represent proteins of a tested protein pair. Haploid double transformants were selected on minimal SD media lacking Leu and Trp. To select for protein-protein interaction, overnight liquid cultures were adjusted to OD_600_=1 and dilution series down to OD_600_=10^-4^ were prepared. 10 µl of the suspensions were dropped onto SD media lacking Leu, Trp and His and containing the appropriate 3-AT concentration and cultivated as described in the previous section.

#### Yeast Three-Hybrid (Y3H) Assays

Genes which code for bridge protein were transferred from pDONR207 into pBRIDGE-GW using Gateway cloning technology. Genes which code for bait proteins were cloned adjacent to Gal4-BD sequence using AQUA cloning method (Beyer et al., 2015). Prey protein constructs were prepared in pACT2-GW vector as mentioned previously. Yeast (*Saccharomyces cerevisiae*) strain Y190 was transformed with pACT2-GW (AD) constructs and pBRIDGE-GW (BD-Bridge) constructs via the lithium acetate (LiAc) method, based on (Gietz and Schiestl, 2007). Co-transformants were selected by cultivation for 2 days on minimal synthetic defined (SD) media (Clontech) lacking Leu (pACT2-GW) or Trp (pBRIDGE-GW). Beta(β)-Galactosidase assay was performed using Yeast β-Galactosidase Assay Kit, Thermo Scientific, with ortho-Nitrophenyl-β-galactoside (ONPG) as substrate. Freshly grown co-transformants in SD-LT and SD-LTM were used in the assay to extract enzyme. Initially OD 600 of the cultures was measured and cells pelleted. Yeast proteins were extracted, and beta-Galactosidase assay solution was added to extract, mixed and incubated for 30 min to 3 hours. Absorbance at 420 nm was measured using the Infinite 200 Pro microplate reader, TECAN. ß-galactosidase activity was calculated using Miller‘s formula, in Miller units (MU) ß-galactosidase activity = (1,000 * Absorbance 420)/(O.D 660* t* V); t= time in minutes of incubation, V = volume of cells used in the assay.

### BTSL1 mutagenesis

Three deletion forms of BTSL1 lacking the region 1137-1142 (RTLVEH) were created: BTSL1-dRH (deletion) and BTSL1-6G (deletion spaced with GGGGGG). An additional deletion form of BTSL1 was lacking the complete M-C site (MSRTLVEHVCREKC). Deletion and 6G substitution were introduced by overlap-extension PCR as described in (Le et al., 2016). Two partially overlapping parts of each sequence were amplified from pDONR207:BTSL1 constructs, using the primer pairs BTSL1_B1 fw/BTSL1_dRH rev, BTSL1_dRH fw/BTSL1_B2 rev (BTSL1-dRH), BTSL1_B1 fw/BTSL1_dRH-G rev, BTSL1_dRH-G fw/BTSL1_B2 rev (BTSL1-6G), BTSL1 B1 fw/ BTSL1_dMC rev, BTSL1_dMC fw/ BTSL1 B2 rev (BTSL1-dMC). Amplicons were transferred into pDONR207, subsequently transferred into pACT2-GW and pGBKT7-GW, transformed into yeast and assayed as described in the previous section.

### Histochemical β-glucuronidase (GUS) Assay

Seedlings were analyzed for GUS activity using 2 mM 5-bromo-4-chloro-3-indoyl-b-D-glucuronic acid (X-Gluc)) as substrate and incubated at 37°C in the dark for 15 min up to 12 h. From *proBTSL1:GUS, proFEP3:GUS* and *proFEP3:FEP3-GUS* lines, four to six seedlings were fixed in ice cold 90% acetone for 1 h and washed in phosphate buffer prior to incubation in the GUS staining solution, which was vacuum infiltrated to obtain better staining. Incubation was performed as described above and stained tissue was fixed in 75% ethanol and 25% acetic acid for 2 h at RT. Chlorophyll was removed by incubation in 70% ethanol for 24 h. Seedlings were imaged with the Axio Imager M2 (Zeiss, 10x objective magnification) and images of entire seedlings assembled with the Stitching function of the ZEN 2 BLUE Edition software (Zeiss).

### Subcellular (co-) localization

To observe subcellular localization, proteins were tagged C-terminally to GFP and/or mCherry fluorophores and/or N-terminally to YFP fluorophore and expressed transiently in tobacco leaf epidermal cells via Agrobacterium-mediated leaf infiltration. For N- and C-terminal fusions, coding sequences were amplified from cDNA of Fe Arabidopsis WT roots with primers carrying B1 and B2 attachment sites, respectively, transferred into the entry vector pDONR207 (Invitrogen) and subcloned into destination vectors pMDC83 (C-terminal GFP fusions) (Curtis and Grossniklaus, 2003), pH7WGY2 (N-terminal YFP) (Karimi et al., 2005), and β-estradiol-inducible pABind-GFP and pABind-mCherry (C-terminal GFP and mCherry, used in co-localization studies) (Bleckmann et al., 2010). The constructs were transformed into Agrobacteria as described in “Plant Material”. A suspension (OD_600_=0.4) of Agrobacteria carrying the construct of interest in infiltration solution (2 mM NaH_2_PO_4_, 0.5% (w/v) glucose, 50 mM MES, 100 µM acetosyringone (in DMSO), pH 5.6) was infiltrated into tobacco leaves using a 1 ml syringe pressed to the abaxial leaf side. For co-localization corresponding Agrobacteria suspensions were mixed 1:1 (each to an OD_600_=0.4) prior to infiltration. For more efficient expression, Agrobacteria carrying the p19 plasmid were co-infiltrated (suppression of RNA interference) (Voinnet et al., 2003, 2015). Transformed plants were kept at RT under long day conditions (16 h light, 8 h dark) and imaged after 48-72 h with a LSM 510 meta confocal laser scanning microscope (Zeiss) or an Axio Imager M2 with ApoTome (Zeiss). GFP and YFP were imaged at an excitation wavelength of 488 nm and emission wavelength of 500 to 530 nm, mCherry was imaged at an excitation wavelength at 563 nm and emission wavelength of 560 to 615 nm. Expression of pABind constructs was induced by spraying β-estradiol mix (20 µM β-estradiol (in DMSO), 0.1% (v/v) Tween20) to the abaxial leaf side 24-48 h post-infiltration (24 h to 48 h before imaging). The (co-) localization experiments were performed in at least two independent replicates or as indicated in the text. Plasmolysis of cells expressing *BTSL1-GFP* was achieved through treatment of the leaf sample with 1 M mannitol solution for 30 min.

### Bimolecular Fluorescence Complementation (BiFC)

CDS of gene pairs to be tested were amplified from cDNA of Arabidopsis WT roots. Amplicons generated with primers carrying B3 and B2 attachment sites were transferred into pDONR221-P3P2 (Thermo Fisher Scientific, for nYFP fusion) and amplicons generated with primers carrying B1 and B4 attachment sites were transferred into pDONR221-P1P4 (Thermo Fisher Scientific, for cYFP fusion), respectively. Primer sequences are listed in **Supplemental Table S1**. In a multisite Gateway LR reaction (Thermo Fisher Scientific), both genes were transferred simultaneously into destination vector pBiFCt-2in1-NN (N-terminal nYFP and cYFP fusions (Grefen and Blatt, 2012), to create pBiFCt-2in1-NN: FEP3:BTSL1, pBiFCt-2in1-NN:PYE-BTSL1, pBiFCt-2in1-NN:PYE-BTSL1-C and pBiFCt-2in1-NN:ILR3-BTSL1-C. The constructs carry a monomeric red fluorescent protein (mRFP) as internal transformation control. As negative controls, structurally similar proteins known to not interact were used (negative controls: pBiFCt-2in1-NN:ILR3-BTSL2-C, pBiFCt-2in1-NN:FIT-BTSL1-C), (Kudla and Bock, 2016). Constructs were transformed into Agrobacteria and subsequently infiltrated into tobacco leaves, as described above. 48 h to 52 h after infiltration, mRFP and YFP signals were detected with an Axio Imager M2 (Zeiss). YFP was imaged at an excitation wavelength of 488 nm and emission wavelength of 500 to 530 nm, mRFP was imaged at an excitation wavelength at 563 nm and emission wavelength of 560 to 615 nm. BiFC experiments were performed in at least two independent replicates with two infiltrated leaves each.

### Gene Expression Analysis by RT-qPCR

Gene expression analysis was performed as described earlier (Abdallah and Bauer, 2016). In brief, mRNA was extracted from whole seedlings grown in the 6 d system (n>60 per replicate) or from roots grown in the 14+3 d system (n>15 per replicate) (see “Plant Growth Conditions”) and used for cDNA synthesis. RT-qPCR was performed using the iTaq™ Universal SYBR® Green Supermix (Bio-Rad) and the SFX96 Touch^TM^ RealTime PCR Detection System (Bio-Rad). Data was processed with the Bio-Rad SFX Manager^TM^ software (version 3.1). Absolute gene expression values were calculated from a gene specific mass standard dilution series and normalized to the elongation factor *EF1Bα*. Primers for mass standards and RT-qPCR are listed in **Supplemental Table S1**. The analysis was performed with three biological and two technical replicates.

### Immunoblot analysis

Total proteins were extracted from ground plant material (tobacco leaves or Arabidopsis whole seedlings grown in the 6 d system, n=30-60 seedlings) with 2x Laemmli buffer (124 mM Tris-HCl, pH 6.8, 5% (w/v) SDS, 4% (w/v) dithithreitol, 20% (v/v) glycerol, with 0.002% (w/v) bromophenol blue) and denatured at 95°C for 10 min. Equal amounts of total protein were separated on SDS-polyacrylamide gels, transferred to a Protran nitrocellulose membrane and stained with PonceauS as described in (Le et al., 2016). To detect HA_3_-tagged FEP3 protein, the membrane was blocked with 5% (w/v) milk solution (Roth) in 1xPBST (137 mM NaCl, 2.7 mM KCl, 10.14 mM Na_2_HPO_4_, 1.76 mM KH_2_PO_4_, 0.1% (v/v) Tween® 20, pH 7.4) for 30 min and subsequently incubated 1 h with anti-HA-peroxidase high-affinity monoclonal rat antibody (3F10; Roche [catalog no. 12013819001]) diluted 1:1000 in 2.5% (w/v) milk solution. After three wash steps, each for 15 min in PBST, the membrane was imaged as described in (Le et al., 2016). Chemiluminescent protein bands were detected with the FluorChem Q system (ProteinSimple) and images were processed with the AlphaView® software (version 3.4.0.0, ProteinSimple).

### Root Length Measurement

Plants were photographed at day six. Length of primary roots of individual seedlings was measured using the JMicroVision software (version 1.2.7, http://www.jmicrovision.com), as described previously (Ivanov et al., 2014). For calculation of mean root lengths and standard deviations, n=13-29 roots per line and condition were measured.

### Seed Fe Content Measurement

To determine seed Fe content, 1-3 plants from each line were grown on soil under long day conditions (16 h light, 8 h dark, 21°C). Seeds were harvested, pooled by plant genotype, and dried for 16 h at 100°C. Fe was extracted from ground seed material by incubation in 500 µl 3% (v/v) HNO_3_ for 16 h at 100°C. Fe content in the supernatant was determined as described (Tamarit et al., 2006). Total Fe content in the sample was calculated with the help of a standard curve and normalized to seed dry weight. Per seed pool, n=3 samples were measured.

### Chlorophyll Content Measurement

Chlorophyll content was measured in leaves of Arabidopsis lines grown in the 10 d system. Chlorophyll a and chlorophyll b were extracted with 100% acetone added to ground leaf material. The supernatant was collected and the washes were repeated until the collected acetone remained colorless. Absorbances of chlorophyll a and b were measured at a wavelength of 662 nm and 645 nm, respectively. Using the absorption coefficients that apply to 100% acetone, chlorophyll a concentration in the measured sample was calculated with c_Chl-a_ [mg µl^-1^]=11.75*A_662_-2.35*A_645_ and chlorophyll b concentration was calculated with c_Chl-b_ [mg µl^-1^]=18.61*A_645_-3.96*A_662_ (Lichtenthaler and Wellburn, 1983), where A is the absorbance at the indicated wavelength in nm. Values were normalized to fresh weight. Four biological replicates were measured, with each biological replicate containing n=5-7 rosettes.

### Multiple Sequence alignments and protein sequence conservation

Multiple sequence alignments were performed with ClustalX using default settings (Larkin et al., 2007). To determine conservation scores of aa in BTSL1, the full BTSL1 aa sequence was uploaded to the Basic Local Alignment Search Tool (BLAST, (Altschul et al., 1990); https://blast.ncbi.nlm.nih.gov/) and run against the Viridiplantae database using the standard blastp (protein-protein BLAST) algorithm. The top 100 hits were downloaded, duplicates were removed. The remaining sequences were used for multiple sequence alignment using the Clustal Omega algorithm (Sievers et al., 2011) and visualized with Jalview (Waterhouse et al., 2009); http://www.jalview.org/). The full aa sequence of FEP3 run against the Viridiplantae database as described above. Hits were only found within the Brassicaceae family, but alignments showed sequence conservation specifically towards the C-terminus. Subsequent blastp of the C-terminal half of FEP3 (25 aa) resulted in several angiosperm hits. FEP3 ortholog sequence hits from exemplary angiosperm orders were downloaded and aligned.

### Protein structure prediction and molecular docking

Protein structures were predicted using AlphaFold2 (Jumper et al., 2021) with protein sequences from TAIR. Multiple Sequence alignments were generated through MMseqs2 API. Molecular docking was performed in HADDOCK 2.4. (van Zundert et al., 2016; Honorato et al., 2021). Active residues were used to generate ambiguous interaction restraints. The obtained file was further processed in Discovery studio, Dassault Systems BIOVIA.

### Statistical Analysis

Null hypothesis between normally distributed groups was tested with a two-tailed Student’s t-test. Null hypothesis was rejected, when the *p*-value (*p*) was below 0.05. Statistically significantly different groups are indicated by one asterisk for *p*<0.05, two asterisks for *p*<0.01 and three asterisks for *p*<0.001. When comparing more than two groups, null hypotheses were tested with one-way analysis of variance (ANOVA) and a Tukey’s post-hoc test. Null hypotheses were rejected when *p*<0.05. Statistically significantly different groups are indicated by different lower-case letters. Number of technical and biological repetitions of the individual experiments are indicated in the respective Methods sections and in the Figure legends.

### Accession Numbers

AKT1 (AT2G26650), BHLH11 (AT4G36060), BHLH34 (AT3G23210), BHLH38 (AT3G56970), BHLH39 (AT3G56980), BHLH100 (AT2G41240), BHLH101 (AT5G04150), BHLH104 (AT4G14410), BHLH115 (AT1G51070), BTS (AT3G18290), BTSL1 (AT1G74770), BTSL2 (AT1G18910), CIPK23 (AT1G30270), DGAT3 (AT1G48300), DUF506 (AT1G12030), FEP1 (AT2G30766), FEP3 (AT1G47400), FIT (AT2G28160), FRO2 (AT1G01580), FRO3 (AT1G23020), GRF11 (AT1G34760), ILR3 (AT5G54680), IRT1 (AT4G19690), JAL12 (AT1G52120), KELCH (AT3G07720), MYB72 (AT1G56160), NAS2 (AT5G56080), NAS4 (AT1G56430), ORG1 (AT5G53450), PRS2 (AT1G32380), PYE (AT3G47640), SDI1 (AT5G48850), S8H (AT3G12900), TCP20 (AT3G27010), UP1 (AT3G06890), UP2 (AT3G56360), UP3 (AT5G05250), URI (AT3G19860),

## Supplemental Material

**Supplemental Figure S1. Targeted Y2H screen (continued in Figure S2).**

**Supplemental Figure S2. Targeted Y2H screen (continued).**

**Supplemental Figure S3. Validation of protein interactions**

**Supplemental Figure S4. Promoter-driven GUS activities of genes encoding the “BTSL-bHLH-FEP3 interactome” in roots**

**Supplemental Figure S5. Subcellular localization of BTSL1-GFP and BTSL1-mCherry (additional images to Figure 3D, YFP-BTSL1).**

**Supplemental Figure S6. FEP3 conserved domain.**

**Supplemental Figure S7. Multiple sequence alignment of bHLH IVc TFs and FEP3 to identify sequence similarities.**

**Supplemental Figure S8. Fe deficiency response phenotypes of FEP3 overexpression lines FEP3-Ox and** *btsl1 btsl2* mutants

**Supplemental Figure S9. Validation of FEP3-OX and *btsl1 btsl2* mutant lines.**

**Supplemental Figure S10. Regulation of selected Fe deficiency response genes in whole seedlings of FEP3-OX and btsl1-btls2.**

**Supplemental Table S1. Primers used in this study.**

## Acknowledgements

We thank Elke Wieneke and Gintaute Matthäi for excellent technical assistance. We thank Ksenia Trofimov for help with imaging of ILR3-GFP and YFP-bHLH104. We acknowledge the contributions of Sarah Plicht, Theresa Priebe, and Kai Blaeser. The authors thank Janneke Balk, Norwich, UK, for *btsl1 btsl2* seeds, and Diqiu Yu, Chinese Academy of Sciences, Kunming, China, for *ProILR3:GUS*/WT and *proBHLH104:GUS*/WT Arabidopsis lines. We thank Ute Hoecker, University of Cologne, Germany, for pBRIDGE plasmid and the Y190 yeast strain, and Andreas Weber, HHU Düsseldorf, Germany, for pGWB3. pACT2-GW and pGBKT7-GW were kindly provided by Dr. Yves Jacob, Institut Pasteur, Paris, France. We thank Rumen Ivanov, Ksenia Trofimov, Inga Mohr, and Tzvetina Brumbarova for help and advice with microscopy and with the implementation of lab protocols. D.M.L and B.S. are members of the international graduate school iGRAD-Plant, Düsseldorf. Funding from the German Research Foundation through the DFG International Research Training groups 1525 and 2466 is greatly acknowledged. This project was funded by the Deutsche Forschungsgemeinschaft (DFG, German Research Foundation) under Germany’s Excellence Strategy – EXC-2048/1 – project ID 390686111. This work was funded by Deutsche Forschungsgemeinschaft GRK F020512056 (NextPlant).

## Abbreviations

3AT: 3-amino-1,2,4-triazole
Aa: Amino acid
AD: Activation domain
BD: Binding domain
bHLH: Basic helix-loop-helix
BiFC: Bimolecular fluorescence complementation
GFP: Green fluorescent protein
GUS: β-Glucuronidase
HHE: Hemerythrin/HHE cation-binding motif
mCherry: Second generation mRFP derivative
mRFP: Monomeric red fluorescent protein
ORF: Open reading frame
OX: Over-expression
RT-qPCR: Reverse transcription quantitative PCR
SD: Standard deviation (Statistics) / Synthetic defined medium (Y2H)
SEP: Small ORF-encoded peptide
TF: Transcription factor
WT: Wild type
Y2H: Yeast two-hybrid
Y3H: Yeast three-hybrid
YFP: Yellow fluorescent protein

## Literature Cited

Aarabi F, Kusajima M, Tohge T, Konishi T, Gigolashvili T, Takamune M, Sasazaki Y, Watanabe M, Nakashita H, Fernie AR, Saito K, Takahashi H, Hubberten HM, Hoefgen R, Maruyama-Nakashita A (2016) Sulfur deficiency-induced repressor proteins optimize glucosinolate biosynthesis in plants. Sci Adv 2: e1601087

Abdallah HB, Bauer P (2016) Quantitative reverse transcription-qPCRbased gene expression analysis in plants. Methods Mol Biol 1363: 9–24

Akmakjian GZ, Riaz N, Guerinot ML (2021) Photoprotection during iron deficiency is mediated by the bHLH transcription factors PYE and ILR3. Proc Natl Acad Sci 118: e2024918118

Altschul SF, Gish W, Miller W, Myers EW, Lipman DJ (1990) Basic local alignment search tool. Journal of Molecular Biology 215: 403–410

Aoki Y, Okamura Y, Tadaka S, Kinoshita K, Obayashi T (2016) ATTED-II in 2016: a plant coexpression database towards lineage-specific coexpression. Plant Cell Physiol 57: e5

Bauer P (2016) Regulation of iron acquisition responses in plant roots by a transcription factor. Biochemistry and Molecular Biology Education 44: 438–449

Bauer P, Ling HQ, Guerinot ML (2007) FIT, the FER-LIKE IRON DEFICIENCY INDUCED TRANSCRIPTION FACTOR in Arabidopsis. Plant Physiol Biochem 45: 260–261

Belda-Palazon B, Julian J, Coego A, Wu Q, Zhang X, Batistic O, Alquraishi SA, Kudla J, An C, Rodriguez PL (2019) ABA inhibits myristoylation and induces shuttling of the RGLG1 E3 ligase to promote nuclear degradation of PP2CA. Plant J

Bensmihen S, To A, Lambert G, Kroj T, Giraudat J, Parcy F (2004) Analysis of an activated ABI5 allele using a new selection method for transgenic Arabidopsis seeds. Febs Letters 561: 127–131

Beyer, H. M., Gonschorek, P., Samodelov, S. L., Meier, M., Weber, W., & Zurbriggen, M. D. (2015). AQUA cloning: a versatile and simple enzyme-free cloning approach. PloS one, 10(9), e0137652.

Bleckmann A, Weidtkamp-Peters S, Seidel CAM, Simon R (2010) Stem cell signaling in Arabidopsis requires CRN to localize CLV2 to the plasma membrane. Plant Physiology 152: 166–176

Briat JF, Curie C, Gaymard F (2007) Iron utilization and metabolism in plants. Current Opinion in Plant Biology 10: 276–282

Brumbarova T, Bauer P, Ivanov R (2015) Molecular mechanisms governing Arabidopsis iron uptake. Trends Plant Sci 20: 124–133

Cabrera-Quio LE, Herberg S, Pauli A (2016) Decoding sORF translation - from small proteins to gene regulation. Rna Biology 13: 1051–1059

Cai Y, Li Y, Liang G (2021) FIT and bHLH Ib transcription factors modulate iron and copper crosstalk in Arabidopsis. Plant, Cell & Environment 44: 1679–1691

Cheng MC, Hsieh EJ, Chen JH, Chen HY, Lin TP (2012) Arabidopsis RGLG2, functioning as a RING E3 ligase, interacts with AtERF53 and negatively regulates the plant drought stress response. Plant Physiology 158: 363–375

Cheng MC, Hsieh EJ, Chen JH, Chen HY, Lin TP (2016) CORRECTION. Arabidopsis RGLG2, Functioning as a RING E3 Ligase, Interacts with AtERF53 and Negatively Regulates the Plant Drought Stress Response. Plant Physiology 170: 1162–1163

Clough SJ, Bent AF (1998) Floral dip: a simplified method for *Agrobacterium*-mediated transformation of *Arabidopsis thaliana*. Plant Journal 16: 735–743

Colangelo EP, Guerinot ML (2004) The essential basic helix-loop-helix protein FIT1 is required for the iron deficiency response. Plant Cell 16: 3400–3412

Curie C, Mari S (2017) New routes for plant iron mining. New Phytologist 214: 521–525

Curtis MD, Grossniklaus U (2003) A gateway cloning vector set for high-throughput functional analysis of genes in planta. Plant Physiology 133: 462–469

Delcourt V, Staskevicius A, Salzet M, Fournier I, Roucou X (2018) Small proteins encoded by unannotated ORFs are rising stars of the proteome, confirming shortcomings in genome annotations and current vision of an mRNA. Proteomics 18

Dong CH, Agarwal M, Zhang Y, Xie Q, Zhu JK (2006) The negative regulator of plant cold responses, HOS1, is a RING E3 ligase that mediates the ubiquitination and degradation of ICE1. Proc Natl Acad Sci U S A 103: 8281–8286

Eide D, Broderius M, Fett J, Guerinot ML (1996) A novel iron-regulated metal transporter from plants identified by functional expression in yeast. Proc Natl Acad Sci U S A 93: 5624–5628

Freemont PS (2000) Ubiquitination: RING for destruction? Curr Biol 10: R84–87

Gao F, Robe K, Bettembourg M, Navarro N, Rofidal V, Santoni V, Gaymard F, Vignols F, Roschzttardtz H, Izquierdo E (2019) The Transcription Factor bHLH121 Interacts with bHLH105 (ILR3) and its Closest Homologs to Regulate Iron Homeostasis in Arabidopsis. The Plant Cell 32: 508–524

Gao F, Robe K, Gaymard F, Izquierdo E, Dubos C (2019) The transcriptional control of iron homeostasis in plants: a tale of bHLH transcription factors? Frontiers in Plant Science 10, https://doi.org/10.3389/fpls.2019.00006

García MJ, Corpas FJ, Lucena C, Alcántara E, Pérez-Vicente R, Zamarreño Á, Bacaicoa E, García-Mina JM, Bauer P, Romera FJ (2018) A shoot Fe signaling pathway requiring the OPT3 transporter controls GSNO reductase and ethylene in *Arabidopsis thaliana* roots. Front Plant Sci 9: 1325

Garcia MJ, Romera FJ, Stacey MG, Stacey G, Villar E, Alcantara E, Perez-Vicente R (2013) Shoot to root communication is necessary to control the expression of iron-acquisition genes in Strategy I plants. Planta 237: 65–75

Gietz RD, Schiestl RH (2007) High-efficiency yeast transformation using the LiAc/SS carrier DNA/PEG method. Nature Protocols 2: 31–34

Graceffa P, Jancsó A, Mabuchi K (1992) Modification of acidic residues normalizes sodium dodecyl sulfate-polyacrylamide gel electrophoresis of caldesmon and other proteins that migrate anomalously. Archives of Biochemistry and Biophysics 297: 46–51

Gratz R, Manishankar P, Ivanov R, Köster P, Mohr I, Trofimov K, Steinhorst L, Meiser J, Mai HJ, Drerup M, Arendt S, Holtkamp M, Karst U, Kudla J, Bauer P, Brumbarova T (2019) CIPK11-dependent phosphorylation modulates FIT activity to promote Arabidopsis iron acquisition in response to calcium signaling. Developmental Cell 48: 726–740.e10

Grefen C, Blatt MR (2012) A 2in1 cloning system enables ratiometric bimolecular fluorescence complementation (rBiFC). Biotechniques 53: 311–314

Grillet L, Lan P, Li WF, Mokkapati G, Schmidt W (2018) IRON MAN is a ubiquitous family of peptides that control iron transport in plants. Nature Plants 4: 953-+

Heim MA, Jakoby M, Werber M, Martin C, Weisshaar B, Bailey PC (2003) The basic helix-loop-helix transcription factor family in plants: a genome-wide study of protein structure and functional diversity. Mol Biol Evol 20: 735–747

Hell R, Stephan UW (2003) Iron uptake, trafficking and homeostasis in plants. Planta 216: 541–551

Hindt MN, Akmakjian GZ, Pivarski KL, Punshon T, Baxter I, Salt DE, Guerinot ML (2017) BRUTUS and its paralogs, BTS LIKE1 and BTS LIKE2, encode important negative regulators of the iron deficiency response in *Arabidopsis thaliana*. Metallomics 9: 876–890

Hirayama T, Lei GJ, Yamaji N, Nakagawa N, Ma JF (2018) The putative peptide gene FEP1 regulates iron deficiency response in Arabidopsis. Plant and Cell Physiology 59: 1739–1752

Honorato RV, Koukos PI, Jiménez-García B, Tsaregorodtsev A, Verlato M, Giachetti A, Rosato A, Bonvin AMJJ (2021) Structural Biology in the Clouds: The WeNMR-EOSC Ecosystem. Frontiers in Molecular Biosciences 8

Hsu PY, Benfey PN (2018) Small but mighty: functional peptides encoded by small ORFs in plants. Proteomics 18: e1700038

Huang TK, Han CL, Lin SI, Chen YJ, Tsai YC, Chen YR, Chen JW, Lin WY, Chen PM, Liu TY, Chen YS, Sun CM, Chiou TJ (2013) Identification of downstream components of ubiquitin-conjugating enzyme PHOSPHATE2 by quantitative membrane proteomics in Arabidopsis roots. Plant Cell 25: 4044–4060

Ivanov R, Brumbarova T, Bauer P (2012) Fitting into the harsh reality: regulation of iron-deficiency responses in dicotyledonous plants. Mol Plant 5: 27–42

Ivanov R, Brumbarova T, Blum A, Jantke AM, Fink-Straube C, Bauer P (2014) SORTING NEXIN1 is required for modulating the trafficking and stability of the Arabidopsis IRON-REGULATED TRANSPORTER1. Plant Cell 26: 1294–1307

Jakoby M, Wang HY, Reidt W, Weisshaar B, Bauer P (2004) FRU (BHLH029) is required for induction of iron mobilization genes in Arabidopsis thaliana. FEBS Lett 577: 528–534

Jefferson RA, Kavanagh TA, Bevan MW (1987) GUS fusions: β-glucuronidase as a sensitive and versatile gene fusion marker in higher plants. EMBO Journal 6: 3901–3907

Jumper J, Evans R, Pritzel A, Green T, Figurnov M, Ronneberger O, Tunyasuvunakool K, Bates R, Žídek A, Potapenko A, Bridgland A, Meyer C, Kohl SAA, Ballard AJ, Cowie A, Romera-Paredes B, Nikolov S, Jain R, Adler J, Back T, Petersen S, Reiman D, Clancy E, Zielinski M, Steinegger M, Pacholska M, Berghammer T, Bodenstein S, Silver D, Vinyals O, Senior AW, Kavukcuoglu K, Kohli P, Hassabis D (2021) Highly accurate protein structure prediction with AlphaFold. Nature 596: 583–589

Kanwar, P., Baby, D., & Bauer, P. (2021). Interconnection of iron and osmotic stress signalling in plants: Is FIT a regulatory hub to cross-connect abscisic acid responses? Plant Biol Suppl 1: 31–38.

Karimi M, De Meyer B, Hilson P (2005) Modular cloning in plant cells. Trends in Plant Science 10: 103–105

Khan MA, Castro-Guerrero NA, McInturf SA, Nguyen NT, Dame AN, Wang J, Bindbeutel RK, Joshi T, Jurisson SS, Nusinow DA, Mendoza-Cozatl DG (2018) Changes in iron availability in Arabidopsis are rapidly sensed in the leaf vasculature and impaired sensing leads to opposite transcriptional programs in leaves and roots. Plant Cell Environ 41: 2263–2276

Kim SA, LaCroix IS, Gerber SA, Guerinot ML (2019) The iron deficiency response in Arabidopsis thaliana requires the phosphorylated transcription factor URI. Proc Natl Acad Sci 116: 24933–24942

Klatte M, Schuler M, Wirtz M, Fink-Straube C, Hell R, Bauer P (2009) The analysis of Arabidopsis nicotianamine synthase mutants reveals functions for nicotianamine in seed iron loading and iron deficiency responses. Plant Physiol 150: 257–271

Kobayashi T (2019) Understanding the complexity of iron sensing and signaling cascades in plants. Plant and Cell Physiology 60: 1440–1446

Kobayashi T, Nagasaka S, Senoura T, Itai RN, Nakanishi H, Nishizawa NK (2013) Iron-binding haemerythrin RING ubiquitin ligases regulate plant iron responses and accumulation. Nat Commun 4: 2792

Kobayashi T, Nozoye T, Nishizawa NK (2018) Iron transport and its regulation in plants. Free Radic Biol Med

Kobayashi T, Nagano AJ, Nishizawa NK (2020) Iron deficiency-inducible peptide-coding genes OsIMA1 and OsIMA2 positively regulate a major pathway of iron uptake and translocation in rice. Journal of Experimental Botany 72: 2196–2211

Koncz C, Schell J (1986) The promoter of T_L_-DNA gene *5* controls the tissue-specific expression of chimeric genes carried by a novel type of Agrobacterium binary vector. Molecular & General Genetics 204: 383–396

Kudla J, Bock R (2016) Lighting the way to protein-protein interactions: recommendations on best practices for Bimolecular Fluorescence Complementation analyses. Plant Cell 28: 1002–1008

Kumar RK, Chu HH, Abundis C, Vasques K, Rodriguez DC, Chia JC, Huang R, Vatamaniuk OK, Walker EL (2017) Iron-nicotianamine transporters are required for proper long distance iron signaling. Plant Physiol 175: 1254–1268

Larkin MA, Blackshields G, Brown NP, Chenna R, McGettigan PA, McWilliam H, Valentin F, Wallace IM, Wilm A, Lopez R, Thompson JD, Gibson TJ, Higgins DG (2007) Clustal W and clustal X version 2.0. Bioinformatics 23: 2947–2948

Le CTT, Brumbarova T, Ivanov R, Stoof C, Weber E, Mohrbacher J, Fink-Straube C, Bauer P (2016) ZINC FINGER OF ARABIDOPSIS THALIANA12 (ZAT12) interacts with FER-LIKE IRON DEFICIENCY-INDUCED TRANSCRIPTION FACTOR (FIT) linking iron deficiency and oxidative stress responses. Plant Physiology 170: 540–557

Lee H, Xiong L, Gong Z, Ishitani M, Stevenson B, Zhu JK (2001) The Arabidopsis HOS1 gene negatively regulates cold signal transduction and encodes a RING finger protein that displays cold-regulated nucleo-cytoplasmic partitioning. Genes & development 15: 912–924

Li X, Zhang H, Ai Q, Liang G, Yu D (2016) Two bHLH transcription factors, bHLH34 and bHLH104, regulate iron homeostasis in *Arabidopsis thaliana*. Plant Physiol 170: 2478–2493

Li Y, Lei R, Pu M, Cai Y, Lu C, Li Z, Liang G (2020) bHLH11 negatively regulates Fe homeostasis by its EAR motifs recruiting corepressors in Arabidopsis. bioRxiv

Li Y, Lu CK, Li CY, Lei RH, Pu MN, Zhao JH, Peng F, Ping HQ, Wang D, Liang G (2021) IRON MAN interacts with BRUTUS to maintain iron homeostasis in *Arabidopsis*. Proceedings of the National Academy of Sciences 118: e2109063118

Liang G, Zhang H, Li X, Ai Q, Yu D (2017) bHLH transcription factor bHLH115 regulates iron homeostasis in *Arabidopsis thaliana*. J Exp Bot 68: 1743–1755

Lichtenthaler HK, Wellburn AR (1983) Determinations of total carotenoids and chlorophylls a and b of leaf extracts in different solvents. Biochemical Society Transactions 11: 591–592

Lindsay WL (1988) Solubility and redox equilibria of iron compounds in soils. In S J.W., G B.A., S U., eds, Iron in Soils and Clay Minerals, Vol 217. Springer, Dordrecht, p 894

Lingam S, Mohrbacher J, Brumbarova T, Potuschak T, Fink-Straube C, Blondet E, Genschik P, Bauer P (2011) Interaction between the bHLH transcription factor FIT and ETHYLENE INSENSITIVE3/ETHYLENE INSENSITIVE3-LIKE1 reveals molecular linkage between the regulation of iron acquisition and ethylene signaling in Arabidopsis. Plant Cell 23: 1815–1829

Liu YG, Mitsukawa N, Oosumi T, Whittier RF (1995) Efficient isolation and mapping of *Arabidopsis thaliana* T-DNA insert junctions by thermal asymmetric interlaced PCR. Plant Journal 8: 457–463

Liu YG, Whittier RF (1995) Thermal Asymmetric Interlaced PCR: automatable amplification and sequencing of insert end fragments from P1 and YAC clones for chromosome walking. Genomics 25: 674–681

Long TA, Tsukagoshi H, Busch W, Lahner B, Salt DE, Benfey PN (2010) The bHLH transcription factor POPEYE regulates response to iron deficiency in Arabidopsis roots. Plant Cell 22: 2219–2236

Mai HJ, Pateyron S, Bauer P (2016) Iron homeostasis in *Arabidopsis thaliana*: transcriptomic analyses reveal novel FIT-regulated genes, iron deficiency marker genes and functional gene networks. BMC Plant Biol 16: 211

Makarewich CA, Olson EN (2017) Mining for Micropeptides. Trends Cell Biol 27: 685–696

Marschner H, Römheld V (1994) Strategies of plants for acquisition of iron. Plant and Soil 165: 261–274

Martín-Barranco A, Spielmann J, Dubeaux G, Vert G, Zelazny E (2020) Dynamic Control of the High-Affinity Iron Uptake Complex in Root Epidermal Cells. Plant Physiol 184: 1236–1250

Matthiadis A, Long TA (2016) Further insight into BRUTUS domain composition and functionality. Plant Signal Behav 11: e1204508

Meiser J, Lingam S, Bauer P (2011) Posttranslational regulation of the iron deficiency basic helix-loop-helix transcription factor FIT is affected by iron and nitric oxide. Plant Physiol 157: 2154–2166

Nakagawa T, Kurose T, Hino T, Tanaka K, Kawamukai M, Niwa Y, Toyooka K, Matsuoka K, Jinbo T, Kimura T (2007) Development of series of gateway binary vectors, pGWBs, for realizing efficient construction of fusion genes for plant transformation. Journal of Bioscience and Bioengineering 104: 34–41

Naranjo-Arcos MA, Maurer F, Meiser J, Pateyron S, Fink-Straube C, Bauer P (2017) Dissection of iron signaling and iron accumulation by overexpression of subgroup Ib bHLH039 protein. Scientific Reports 7, 10911

Nouet C, Motte P, Hanikenne M (2011) Chloroplastic and mitochondrial metal homeostasis. Trends Plant Sci 16: 395–404

Ohkubo Y, Tanaka M, Tabata R, Ogawa-Ohnishi M, Matsubayashi Y (2017) Shoot-to-root mobile polypeptides involved in systemic regulation of nitrogen acquisition. Nat Plants 3: 17029

Palmer CM, Hindt MN, Schmidt H, Clemens S, Guerinot ML (2013) MYB10 and MYB72 are required for growth under iron-limiting conditions. PLoS Genet 9: e1003953

Robinson NJ, Procter CM, Connolly EL, Guerinot ML (1999) A ferric-chelate reductase for iron uptake from soils. Nature 397: 694–697

Rodriguez-Celma J, Chou H, Kobayashi T, Long TA, Balk J (2019) Hemerythrin E3 Ubiquitin Ligases as Negative Regulators of Iron Homeostasis in Plants. Front Plant Sci 10: 98

Rodríguez-Celma J, Connorton JM, Kruse I, Green RT, Franceschetti M, Chen Y-T, Cui Y, Ling H-Q, Yeh K-C, Balk J (2019) Arabidopsis BRUTUS-LIKE E3 ligases negatively regulate iron uptake by targeting transcription factor FIT for recycling. Proceedings of the National Academy of Sciences: 201907971

Rodríguez-Celma J, Green RT, Connorton JM, Kruse I, Cui Y, Ling HQ, Balk J (2017) BRUTUS-LIKE proteins moderate the transcriptional response to iron deficiency in roots. bioRxiv [Preprint]: doi.org/10.1101/231365

Salahudeen AA, Thompson JW, Ruiz JC, Ma HW, Kinch LN, Li Q, Grishin NV, Bruick RK (2009) An E3 ligase possessing an iron-responsive hemerythrin domain is a regulator of iron homeostasis. Science 326: 722–726

Samira R, Li BH, Kliebenstein D, Li CY, Davis E, Gillikin JW, Long TA (2018) The bHLH transcription factor ILR3 modulates multiple stress responses in Arabidopsis. Plant Molecular Biology 97: 297–309

Schmid NB, Giehl RF, Döll S, Mock HP, Strehmel N, Scheel D, Kong X, Hider RC, von Wirén N (2014) Feruloyl-CoA 6’-Hydroxylase1-dependent coumarins mediate iron acquisition from alkaline substrates in Arabidopsis. Plant Physiol 164: 160–172

Schuler M (2011) The role of nicotianamine in the metal homeostasis of *Arabidopsis thaliana*. Saarländische Universitäts- und Landesbibliothek [Dissertation]: doi:10.22028/D22291-22738

Schuler M, Rellán-Álvarez R, Fink-Straube C, Abadía J, Bauer P (2012) Nicotianamine functions in the phloem-based transport of iron to sink organs, in pollen development and pollen tube growth in Arabidopsis. Plant Cell 24: 2380–2400

Schwarz B, Bauer P (2020) FIT, a regulatory hub for iron deficiency and stress signaling in roots, and FIT-dependent and-independent gene signatures. Journal of Experimental Botany

Selote D, Samira R, Matthiadis A, Gillikin JW, Long TA (2015) Iron-binding E3 ligase mediates iron response in plants by targeting basic helix-loop-helix transcription factors. Plant Physiol 167: 273–286

Sievers F, Wilm A, Dineen D, Gibson TJ, Karplus K, Li WZ, Lopez R, McWilliam H, Remmert M, Söding J, Thompson JD, Higgins DG (2011) Fast, scalable generation of high-quality protein multiple sequence alignments using Clustal Omega. Molecular Systems Biology 7

Sivitz A, Grinvalds C, Barberon M, Curie C, Vert G (2011) Proteasome-mediated turnover of the transcriptional activator FIT is required for plant iron-deficiency responses. Plant Journal 66: 1044–1052

Sivitz AB, Hermand V, Curie C, Vert G (2012) Arabidopsis bHLH100 and bHLH101 control iron homeostasis via a FIT-independent pathway. PLoS One 7: e44843

Stührwohldt N, Schaller A (2019) Regulation of plant peptide hormones and growth factors by post-translational modification. Plant Biol (Stuttg) 21 Suppl 1: 49–63

Tabata R, Sumida K, Yoshii T, Ohyama K, Shinohara H, Matsubayashi Y (2014) Perception of root-derived peptides by shoot LRR-RKs mediates systemic N-demand signaling. Science 346: 343–346

Tamarit J, Irazusta V, Moreno-Cermeño A, Ros J (2006) Colorimetric assay for the quantitation of iron in yeast. Analytical Biochemistry 351: 149–151

Tanabe N, Noshi M, Mori D, Nozawa K, Tamoi M, Shigeoka S (2019) The basic helix-loop-helix transcription factor, bHLH11 functions in the iron-uptake system in Arabidopsis thaliana. Journal of plant research 132: 93–105

Tirode, F., Malaguti, C., Romero, F., Attar, R., Camonis, J., & Egly, J. M. (1997). A conditionally expressed third partner stabilizes or prevents the formation of a transcriptional activator in a three-hybrid system. Journal of Biological Chemistry, 272(37), 22995–22999.

Tissot N, Robe K, Gao F, Grant-Grant S, Boucherez J, Bellegarde F, Maghiaoui A, Marcelin R, Izquierdo E, Benhamed M, Martin A, Vignols F, Roschzttardtz H, Gaymard F, Briat JF, Dubos C (2019) Transcriptional integration of the responses to iron availability in Arabidopsis by the bHLH factor ILR3. New Phytol

Tottey S, Block MA, Allen M, Westergren T, Albrieux C, Scheller HV, Merchant S, Jensen PE (2003) Arabidopsis CHL27, located in both envelope and thylakoid membranes, is required for the synthesis of protochlorophyllide. Proc Natl Acad Sci U S A 100: 16119–16124

Tsugeki R, Kochieva EZ, Fedoroff NV (1996) A transposon insertion in the Arabidopsis SSR16 gene causes an embryo-defective lethal mutation. Plant Journal 10: 479–489

van Zundert GCP, Rodrigues JPGLM, Trellet M, Schmitz C, Kastritis PL, Karaca E, Melquiond ASJ, van Dijk M, de Vries SJ, Bonvin AMJJ (2016) The HADDOCK2.2 Web Server: User-Friendly Integrative Modeling of Biomolecular Complexes. Journal of Molecular Biology 428: 720–725

Vert G, Grotz N, Dedaldechamp F, Gaymard F, Guerinot ML, Briat JF, Curie C (2002) IRT1, an Arabidopsis transporter essential for iron uptake from the soil and for plant growth. Plant Cell 14: 1223–1233

Vert GA, Briat JF, Curie C (2003) Dual regulation of the Arabidopsis high-affinity root iron uptake system by local and long-distance signals. Plant Physiology 132: 796–804

Voinnet O, Rivas S, Mestre P, Baulcombe D (2003) An enhanced transient expression system in plants based on suppression of gene silencing by the p19 protein of tomato bushy stunt virus (Retracted article. See vol. 84, pg. 846, 2015). Plant Journal 33: 949–956

Voinnet O, Rivas S, Mestre P, Baulcombe D (2015) An enhanced transient expression system in plants based on suppression of gene silencing by the p19 protein of tomato bushy stunt virus (Retraction of Vol 33, Pg 949, 2003). Plant Journal 84: 846–846

von Wirén N, Klair S, Bansal S, Briat JF, Khodr H, Shioiri T, Leigh RA, Hider RC (1999) Nicotianamine chelates both Fe-III and Fe-II. Implications for metal transport in plants. Plant Physiology 119: 1107–1114

Wang HY, Klatte M, Jakoby M, Bäumlein H, Weisshaar B, Bauer P (2007) Iron deficiency-mediated stress regulation of four subgroup Ib BHLH genes in *Arabidopsis thaliana*. Planta 226: 897–908

Wang N, Cui Y, Liu Y, Fan H, Du J, Huang Z, Yuan Y, Wu H, Ling HQ (2013) Requirement and functional redundancy of Ib subgroup bHLH proteins for iron deficiency responses and uptake in *Arabidopsis thaliana*. Mol Plant 6: 503–513

Waterhouse AM, Procter JB, Martin DMA, Clamp M, Barton GJ (2009) Jalview Version 2-a multiple sequence alignment editor and analysis workbench. Bioinformatics 25: 1189–1191

Wedepohl KH (1995) The composition of the continental crust. Geochimica Et Cosmochimica Acta 59: 1217–1232

Xu J, Li HD, Chen LQ, Wang Y, Liu LL, He L, Wu WH (2006) A protein kinase, interacting with two calcineurin B-like proteins, regulates K^+^ transporter AKT1 in Arabidopsis. Cell 125: 1347–1360

Yang JL, Chen WW, Chen LQ, Qin C, Jin CW, Shi YZ, Zheng SJ (2013) The 14-3-3 protein GENERAL REGULATORY FACTOR11 (GRF11) acts downstream of nitric oxide to regulate iron acquisition in *Arabidopsis thaliana*. New Phytol 197: 815–824

Young JM, Kuykendall LD, Martinez-Romero E, Kerr A, Sawada H (2001) A revision of Rhizobium Frank 1889, with an emended description of the genus, and the inclusion of all species of *Agrobacterium* Conn 1942 and *Allorhizobium undicola* de Lajudie *et al*. 1998 as new combinations: *Rhizobium radiobacter*, *R. rhizogenes*, *R. rubi*, *R. undicola* and *R. vitis*. International Journal of Systematic and Evolutionary Microbiology 51: 89–103

Yuan Y, Wu H, Wang N, Li J, Zhao W, Du J, Wang D, Ling HQ (2008) FIT interacts with AtbHLH38 and AtbHLH39 in regulating iron uptake gene expression for iron homeostasis in Arabidopsis. Cell Res 18: 385–397

Zanet J, Benrabah E, Li T, Pélissier-Monier A, Chanut-Delalande H, Ronsin B, Bellen HJ, Payre F, Plaza S (2015) Pri sORF peptides induce selective proteasome-mediated protein processing. Science 349: 1356–1358

Zhai Z, Gayomba SR, Jung HI, Vimalakumari NK, Piñeros M, Craft E, Rutzke MA, Danku J, Lahner B, Punshon T, Guerinot ML, Salt DE, Kochian LV, Vatamaniuk OK (2014) OPT3 is a phloem-specific iron transporter that is essential for systemic iron signaling and redistribution of iron and cadmium in Arabidopsis. Plant Cell 26: 2249–2264

Zhang J, Liu B, Li M, Feng D, Jin H, Wang P, Liu J, Xiong F, Wang J, Wang HB (2015) The bHLH transcription factor bHLH104 interacts with IAA-LEUCINE RESISTANT3 and modulates iron homeostasis in Arabidopsis. Plant Cell 27: 787–805

